# Individual variation in dispersal, and its sources, shape the fate of pushed vs. pulled range expansions

**DOI:** 10.1101/2022.01.12.476009

**Authors:** Maxime Dahirel, Chloé Guicharnaud, Elodie Vercken

**Affiliations:** Université Côte d’Azur, INRAE, CNRS, ISA, France; Ghent University, Department of Biology, Belgium

**Keywords:** density-dependent dispersal, dispersal evolution, individual-based models, expansion velocity, genetic diversity, heritability

## Abstract

Ecological and evolutionary dynamics of range expansions are shaped by both dispersal and population growth. Accordingly, density-dependence in either dispersal or growth can determine whether expansions are pulled or pushed, i.e. whether expansion velocities and genetic diversity are mainly driven by recent, low-density edge populations, or by older populations closer to the core. Despite this and despite abundant evidence of dispersal evolution during expansions, the impact of density-dependent dispersal and its evolution on expansion dynamics remains understudied. Here, we used simulation models to examine the influence of individual trait variation in both dispersal capacity and dispersal density- dependence on expansions, and how it impacts the position of expansions on the pulled-pushed continuum. First, we found that knowing about the evolution of density-dependent dispersal at the range edge can greatly improve our ability to predict whether an expansion is (more) pushed or (more) pulled. Second, we found that both dispersal costs and the sources of variation in dispersal (genetic or non-genetic, in dispersal capacity versus in density- dependence) greatly influence how expansion dynamics evolve. Among other scenarios, pushed expansions tended to become more pulled with time only when density-dependence was highly heritable, dispersal costs were low and dispersal capacity could not evolve. When, on the other hand, variation in density-dependence had no genetic basis, but dispersal capacity could evolve, then pushed expansions tended to become more pushed with time, and pulled expansions more pulled. More generally, our results show that trying to predict expansion velocities and dynamics using trait information from non-expanding regions only may be problematic, that both dispersal variation and its sources play a key role in determining whether an expansion is and stays pushed, and that environmental context (here dispersal costs) cannot be neglected. Those simulations suggest new avenues of research to explore, both in terms of theoretical studies and regarding ways to empirically study pushed vs. pulled range expansions.

## Introduction

By redistributing both genes and individuals in space, often non-randomly, dispersal has the potential to influence many aspects of ecological dynamics (e.g. Jacob et al., 2019; Little et al., 2019). Understanding what drives dispersal is especially important in the context of range expansions and climate-driven range shifts, as spread in space is the product of dispersal and population growth dynamics (Lewis et al., 2016). Dispersal is a complex trait with multiple and interacting drivers (Bowler & Benton, 2005; Matthysen, 2012). Intuitively, one may assume that qualitatively different dispersal or growth functions would therefore lead to qualitatively different expansion dynamics. In the context of range expansions, this has been especially studied with respect to density-dependence. One can indeed differentiate between “pushed” or “pulled” expansions (or maybe more accurately place them on a “pushiness” gradient), based on the density-dependence of dispersal and/or growth (Birzu et al., 2019; Lewis et al., 2016; Stokes, 1976). Pulled expansions are predicted to happen when the product of dispersal and growth is maximal at low densities; spread is then primarily driven by what happens in the low-density edge populations where dispersal or growth are maximal. On the other side of the continuum, pushed expansions should happen when dispersal and/or growth are instead increasing with population density, due to e.g. density-dependent dispersal or Allee effects (Birzu et al., 2019). In that case, the contributions of dispersal and growth at the low-density edge to overall spread are outweighed by what happens in older more populated habitats; expansions are thus “pushed” forward by these older populations. The more pushed an expansion is, the more predictions of spread velocity based only on low-density behaviour will underestimate its true speed (Birzu et al., 2018, 2019; Gandhi et al., 2016, 2019)

Evolutionary dynamics during expansions are likely influenced by whether they are pushed or pulled (Miller et al., 2020): pulled expansions are typically predicted to lose genetic diversity faster than pushed ones, as new populations are founded by fewer individuals (Roques et al., 2012). On the one side, higher genetic diversity in pushed expansions may mean they are better equipped to successfully adapt to new environmental conditions encountered during expansion (Szűcs et al., 2017). On the other hand, the fact that founding population sizes may be larger at the edge in pushed expansions may mean that spatial sorting processes (Phillips & Perkins, 2019 and see below) are less selective (Miller et al., 2020). In contrast to this, the traditional view of the pushed/pulled expansion distinction in theoretical works implicitly assumes there is no individual variation, and thus no evolution, in the very traits that generate “pushiness”; that is, the density-dispersal and density-growth functions are the same for all individuals, and remain constant during the expansion.

Individual variation in dispersal and dispersal-related traits is very common and now well documented (Bowler & Benton, 2005; Cote et al., 2017; Jacob et al., 2019; Ronce & Clobert, 2012), and while a lot of questions remain, it is clear that a non-negligible fraction of this variation is genetic (Saastamoinen et al., 2018). Following Cote et al. (2017)’s terminology, we can distinguish individual variation in dispersal capacity/ability, which is linked to variation in enabling traits, making dispersal possible at all (e.g. presence/ absence of wings, Simmons & Thomas, 2004) or enhancing traits (e.g. body condition or wing/leg length, Baines et al., 2019; Baines & McCauley, 2018), from individual variation in context-dependence, which can be linked to matching traits that do not alter dispersal capacity *per se*, but lead to non-random dispersal in response to experienced environmental conditions.

We also know that this individual variation can influence spread dynamics (Miller et al., 2020). Genetic differences in dispersal traits can fuel evolution at the expanding range edge by spatial sorting, as at each generation, individuals with spread-facilitating traits are more likely to advance to new habitats and reproduce there (Phillips & Perkins, 2019; Shine et al., 2011). The effects of spatial sorting on spread velocity are well documented, thanks to both iconic natural examples (e.g. Phillips et al., 2006) and experiments reshuffling individuals to cancel out the effects of spatial evolution (Ochocki & Miller, 2017; Weiss-Lehman et al., 2017). Because dispersal is often associated with other traits in syndromes (Ronce & Clobert, 2012), changes in many traits along expansions are potentially linked with spatial sorting (Chuang & Peterson, 2016).

However, these studies have so far mostly focused on variation in dispersal capacity (or the underlying enabling or enhancing traits). As a result, and despite some indications that individual variation in density-dependence itself may be subject to selection during range expansions (Dahirel, Bertin, Calcagno, et al., 2021; Fronhofer et al., 2017; Mishra et al., 2020; Weiss-Lehman et al., 2017), we mostly don’t know how the structure of individual variation in dispersal influences whether an expansion is and stays pushed or pulled. Indeed, the one theoretical study on the subject, which showed that genetic variation leads pushed expansions to become pulled with time, was focused on Allee effects only (Erm & Phillips, 2020). In the context of dispersal, theory developed outside of the pushed/pulled framework suggests that dispersal can become more density-independent at range edges (Travis et al., 2009), which would agree with Erm and Phillips (2020)’s predictions. However, that theoretical study and others typically allow only positive density-dependent dispersal to evolve, when negative density-dependent dispersal is just as likely in nature (Harman et al., 2020). This may explain mismatches with some of the few existing experimental studies (Dahirel, Bertin, Calcagno, et al., 2021; Fronhofer et al., 2017; Mishra et al., 2020). More importantly, the fact that dispersal becomes *on average* more density-independent across the range of densities tells us nothing in itself about the difference between average/maximal dispersal and specifically low-density dispersal. It is that difference which should *a priori* matter for the position of an expansion on the pushed/pulled gradient.

In this context, we used individual-based simulations to examine how genetic and non-genetic sources of inter-individual trait variation influence the evolution and maintenance of pushed vs pulled expansion dynamics. We focused on dispersal variation, rather than on Allee effects as many studies on pushed expansions did so far (e.g. Birzu et al., 2018; Erm & Phillips, 2020; Gandhi et al., 2016; Roques et al., 2012; but see Birzu et al., 2019) for two reasons. First, while there are reasons to believe the broad qualitative results hold whether pushed expansions are caused by Allee effects or dispersal (Birzu et al., 2019), this still needs to be confirmed, especially as evolution of dispersal-density reaction norms would interact with selection for higher dispersal at range edges. Second, although there are no directly comparable systematic syntheses, available evidence suggests positive density-dependent dispersal is more frequent than (demographic) Allee effects in nature (Gregory et al., 2010; Harman et al., 2020), something that is not (yet) reflected in the literature on pushed expansions. We examined how dispersal influences whether expansions are pushed or pulled by drawing on several lines of evidence, including genetic diversity information (Gandhi et al., 2016, 2019). We determined how the sources of dispersal variation (genetic vs. non genetic, in dispersal capacity or dispersal matching traits), influence the evolution of “pushiness”. Finally, we analyzed how dispersal mortality interact with these sources of variation to shape dispersal evolution during range expansions. In non-expanding populations at least, higher costs are predicted to favour more positive density-dependent dispersal (Rodrigues & Johnstone, 2014; Travis et al., 1999), which would influence the evolution of pushed vs. pulled dynamics.

## Methods

Our discrete-time and discrete-space simulation model was written in NetLogo (Wilensky, 1999) version 6.2.0. We interacted with the model and designed our experiments using the nlrx R package (Salecker et al., 2019). We provide below a summarised description of the model, before presenting the simulation experiments we ran using it. A detailed description of the simulation model using the ODD protocol (Grimm et al., 2010; Grimm et al., 2020) can found in **Supplementary Material S1**, and a copy is attached with the model and analysis code (**Data availability**).

### Summary model description

The overall purpose of our model was to understand how dispersal trait distribution and dispersal evolution shape evolutionary dynamics during range expansions, with a focus on density-dependence and the emergence of pushed vs pulled dynamics. Briefly, the model describes the dynamics of asexual haploid individuals spreading in a linear discrete landscape, with their growth dynamics shaped by a Ricker model and their dispersal by Kun and Scheuring (2006)’s model (**Fig. 1**). Dispersal and growth are both stochastic. Individuals can vary or not in traits shaping dispersal, and that variation can be more or less heritable. Dispersal is limited to the nearest neighbours, and dispersal costs can be imposed in the form of a mortality risk.

**Figure 1.**
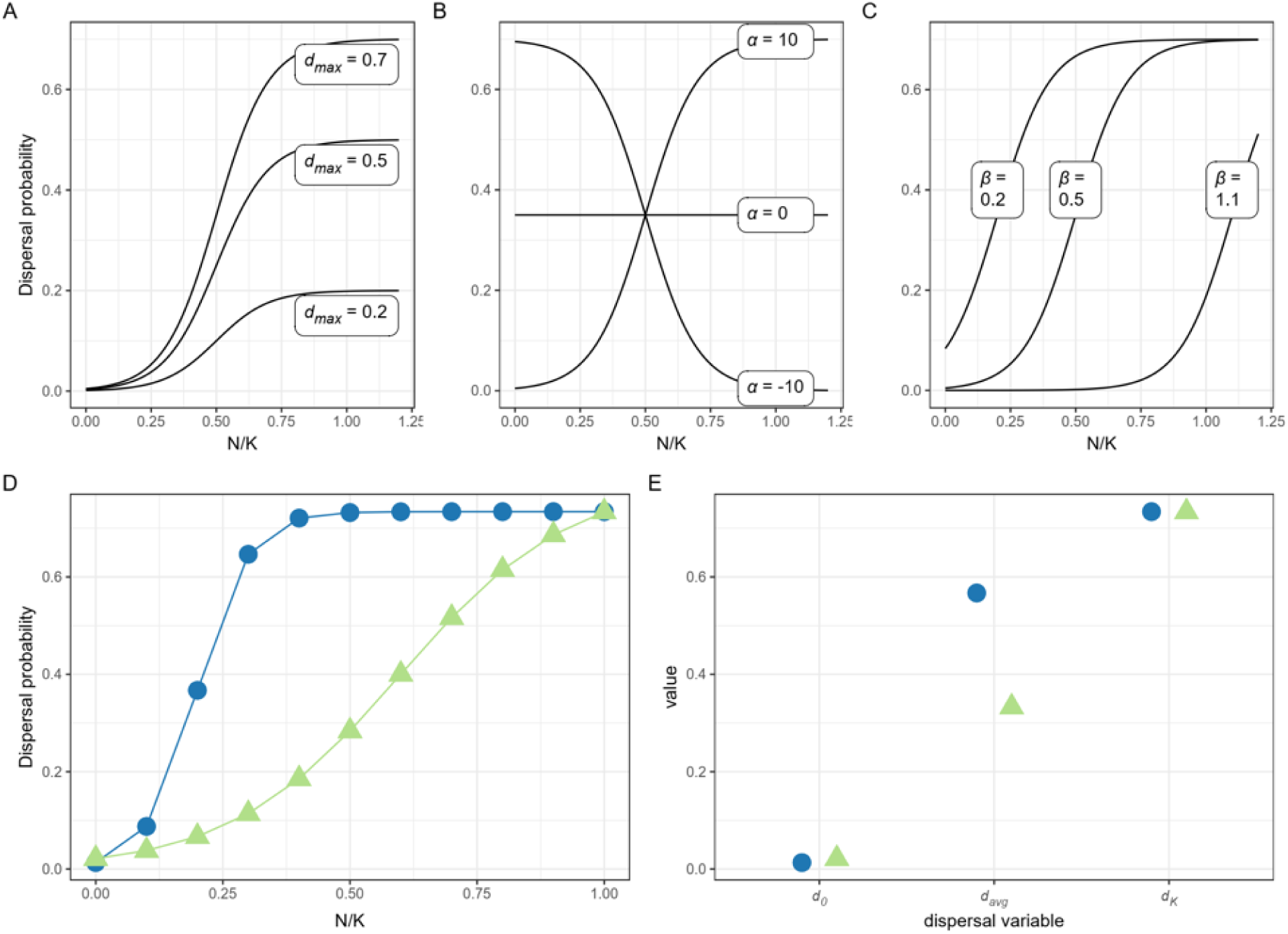
Illustration of the dispersal function used in the individual-based model (based on Kun & Scheuring, 2006) and some of its properties. A, B, C: effect of changing one of the three underlying variables while keeping the other two constant. Values when kept constant: *d*_*max*_=0.5, *α* = 10, *β* =0.5. D: illustration of the sampling points used to estimate *d*_*avg*_ for two example functions, E: effect of the function general shape on *d*_*avg*_. Note how two functions can have (nearly) the same *d*_0_ and *d*_*K*_ (and therefore the same density-dependence *Δ*_*K*−0_), while differing in *d*_*avg*_ (and therefore in *Δ*_*avg*−0_).

The model world in which our virtual species expands is one-dimensional, has closed boundaries, and is made of discrete patches in which individuals live and between which they may disperse. Patches, which form the spatial units of the model, are characterized by their location *x*(integer ≥0), by their current population size *N*, and by their carrying capacity *K*, the latter being equal and constant among all patches.

Individuals each possess three dispersal traits, the maximal dispersal rate *d*_*max*_, the slope *α*, and the midpoint *β*. Individual dispersal probability then depends on these traits and local population density *N* at the start of the dispersal phase (**Fig. 1**, Kun & Scheuring, 2006). Following the terminology in Cote et al (2017), variation in *d*_*max*_ can be interpreted as encoding variation in dispersal enhancing or enabling traits, while variation in *α* or *β* corresponds to variation in dispersal matching traits. We allow *β* to be negative instead of being constrained to be always ≥0, as this is the most convenient way to generate flat dispersal-density functions (see detailed discussion in **Supplementary Materials S1 and S2**). To allow for genetic and non-genetic variation, each trait is actually the (transformed) sum of an underlying additive genetic component (logit(*d*_*max*_)_[*a*]_, *α*_[*a*]_, *β*_[*a*]_) and an underlying noise/residual part (logit(*d*_*max*_)_[*r*]_, *α*_[*r*]_, *β*_[*r*]_). To track neutral genetic diversity, individuals also possess a neutral locus *γ* at which two allelic values are possible (0 or 1).

Once we know the *d*_*max*_, *α* and *β* of an individual, we additionally calculate five values at the individual level (**Fig. 1D-E**):

- *d*_0_, the hypothetical dispersal rate at *N* =0. The theoretical distinction between pulled and pushed expansions hinges on whether expansions move as fast as expected from *d*_0_ or faster (e.g. Birzu et al., 2019);
- *d*_*K*_, the expected dispersal rate at *N* = *K*;
- *d*_*avg*_, the average dispersal over the range of densities 0 − *K*. For computational reasons, we approximate it by calculating *d*_*N*_ every 0.1*K* from 0 to *K*;
- two measures of the strength of density-dependence *Δ*_*K*−0_ = *d*_*K*_ − *d*_0_ and *Δ*_*avg*−0_ = *d*_*avg*_ − *d*_0_. The former is independent of the shape of the dispersal function between 0 and *K*, while the latter accounts for it at least in part.

The model is initialised by releasing *K* adult individuals that have not yet dispersed nor reproduced in the patch *x*=0. Initial values for the genetic and non-genetic components of individual traits are drawn from Normal distributions, with the value of initial heritability *h*^2^ used to determine how initial phenotypic variance is partitioned between genetic and non-genetic components (see **Virtual experiment design** below and **Table 1** for the means and variances). Initial allelic values for the neutral locus are drawn from Bernoulli(0.5). The world is set to be long enough that the expansion never runs out of patches. As we assume stepping- stone dispersal (dispersers only move one patch) and the expansion advances in one direction, this means, in practice, that the world can be any number of patches as long as it is larger than the number of generations in a run (for run duration in our experiment, see **Virtual experiment design**).

**Table 1.** Summary of global parameters used in our simulation model, and their possible values.

| Parameter name | Description | Initial values |
| --- | --- | --- |
| $m$ | dispersal mortality | 0.1; 0.5; 0.9 |
| $h^2$ | initial trait heritability | 0; 0.33; 1 |
| $r_0$ | mean low-density growth rate | $\log(5)$ |
| $K$ | patch carrying capacity | 250 |
| $\overline{d_{max}}_{t=0}$ | initial average $d_{max}$ | 0.5 |
| $\overline{\alpha}_{t=0}$ | initial average relative slope parameter of the dispersal-density function | -6; 0; 6 |
| $\overline{\beta}_{t=0}$ | initial average midpoint of the dispersal-density function | 0; 0.5; 1 |
| $V_{P[\logit(d_{max})]}$ | initial phenotypic variance for $d_{max}$ (on the logit scale) | 0; $1.5^2$ |
| $(V_{P[\alpha]}, V_{P[\beta]})$ | vector of initial phenotypic variances for $\alpha$ and $\beta$ | (0, 0) ; ($6^2$, $0.5^2$) |

Generations are non-overlapping, and every time step, the life cycle unfolds as follows:

- Individuals are counted, providing information about patch population sizes *N* for dispersal;
- *Dispersal:* Newly adult individuals may disperse with a probability *d* depending on their individual traits (*d_max_, α, β*) and local population size 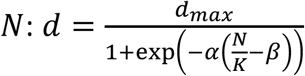 (**Fig. 1**, Kun & Scheuring, 2006). Individuals that do disperse die with a probability *m*; if they survive, they settle in one of the immediate neighbouring patches, randomly chosen.
- Individuals are re-counted post-dispersal, updating population sizes *N* for the reproduction phase;
- *Reproduction:* Each remaining adult then produces *F* juveniles, with *F* ∼ Poisson(*λ*) and the mean fecundity *λ* based on a Ricker model: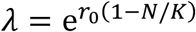, where *r*_0_ is the low-density growth rate. Juveniles are born in the patch currently occupied by their parent, and directly inherit their genetic values for dispersal-related and neutral loci. Their values for the noise part of the dispersal traits are redrawn at random.
- *Death and end of cycle:* All adults die; juveniles then become adults.

To facilitate further analyses, a series of patch-level summaries of traits are computed every generation. In this study, we are particularly interested in the means of *d*_*max*_, *d*_0_, the density-dependence metrics *Δ*_*K*−0_, *Δ*_*avg*−0_, and the neutral allele frequencies (all collected right before the dispersal phase).

### Simulation experiments and analyses

#### Virtual experiment design

We tested all possible combinations of the starting parameters described in **Table 1**, with the exception of the effectively redundant ones (namely heritabilities *h*^2^ >0 when all phenotypic variances were 0), resulting in 270 scenarios. We ran expansions for 120 generations, and “observed” them every 20 generations to save memory. We replicated each scenario 10 times. Besides the extreme *h*^2^ =0 and *h*^2^ = 1 cases, the “intermediate” level of initial heritability was set to 0.33 to roughly match “average” heritabilities seen in dispersal traits (Saastamoinen et al., 2018). The growth rate *r*_0_ was constant and set to a value that is within the range of plausible values for insects (Hassell et al., 1976), and worked well in previous simulations (Dahirel, Bertin, Haond, et al., 2021). *K* was chosen to match previous simulations (Dahirel, Bertin, Haond, et al., 2021; Haond et al., 2018), and is within an order of magnitude of typical population sizes seen during experimental range expansions in plants or invertebrates (e.g. Dahirel, Bertin, Haond, et al., 2021; Ochocki & Miller, 2017; Van Petegem et al., 2018; Williams & Levine, 2018). The value for the initial phenotypic variance for *logit*(*d*_*max*_) was chosen because a Normal(*μ* =0, *σ* = 1.5) distribution on the logit scale leads to an approximately uniform distribution over most of the [0;1] range on the proportion scale. The non-zero starting values for the initial mean slope *α* were chosen based on a key property of the logistic function: *d* goes from 0.05 × *d*_*max*_to 0.95 × *d*_*max*_over an interval of width ≃ 6/|*α*| centered on *β* (see e.g. Börger & Fryxell, 2012). Setting initial |*α*| to be 6 means that interval is of width *K*. The initial phenotypic variance for *α* was chosen so that the mean initial difference between replicates with *α* =0 and replicates with *α* ≠0 was 1 standard deviation; a similar argument was used to set the initial phenotypic variance for the midpoint *β*.

We repeated the entire experimental design and analyses with a lower growth rate (*r*_0_ = log(1.5)) to determine whether our results on dispersal evolution were sensitive to overall fecundity. Most of our key results (i.e. results pertaining to density-dependence) are upheld; deviations to the results observed when *r*_0_ = log(5) are briefly described in **Results** and **Discussion**.

#### Analyses

All analyses were done using R, version 4.1.2 (R Core Team, 2021). Simulated data handling, analyses and plotting relied mostly on the `mgcv` (Wood, 2017), `gratia` (Simpson, 2021), ordinal (Christensen, 2019), `MuMIn` (Barton, 2020) and `patchwork` (Pedersen, 2020) packages, as well as the `tidyverse` suite of packages (Wickham et al., 2019).

We are interested here in how traits evolve at the range front, i.e. the area at the edge of the expansion where a density gradient is present (e.g. Lewis et al., 2016), and how this influences expansion dynamics. This means we first need to define what patches belong to the range front. Based on plots showing the distribution of population sizes as a function of distance to the farthest-forward population (**Supplementary Material S3**), we decided all patches less than 5 patches from the limit of the expansion belong to the range front for our purposes. Patches closer to the core typically have already reached an equilibrium density (which may or may not be *K*, depending on the interplay between growth and dispersal in the core; **Supplementary Material S3**). For each replicate expansion, we used weighted averages (based on population size at the time of trait measurement) to average trait values across front patches before analyses. We ran separate sets of statistical models on runs with *r*_0_ = log(5) and runs with *r*_0_ = log(1.5).

### Expansion velocity

Broadly speaking, and unless *r*_0_ is negative (strong Allee effect, Courchamp et al., 2008), theory predicts that the long run velocity *v* of an expansion can be described as *v* = *p* × *v*_*F*_ = *p* × *f*(*d*_0_, *r*_0_), where *v*_*F*_ is the velocity expected for a pulled expansion with these *r*_0_ and *d*_0_ (Birzu et al., 2018, 2019; Gandhi et al., 2016; Lewis et al., 2016; Wang et al., 2019). What we term *p* here is a measure of “pushiness”, and increases the more growth or dispersal are positively density dependent (Birzu et al., 2018, 2019). Although analytical formulas for *v*_*F*_ exist in many situations (Wang et al., 2019), they are mostly designed for cases when population densities are high and continuous, rather than counts of discrete individuals. While the resulting bias may be small to negligible in some scenarios (Dahirel, Bertin, Haond, et al., 2021; Haond et al., 2018), the approximations can become important in some cases, for instance when stochasticity is high (e.g. Hallatschek & Korolev, 2009), and available corrections are not always practical to implement.

To circumvent these issues and clearly separate the effect of *d*_0_ from the effect of density-dependence, we decided instead to analyse expansion velocities using Generalized Additive Models (GAMs). This allowed us to describe expansion velocities using very flexible non-linear functions of dispersal traits (growth rates being identical across all replicates), without having to choose ourselves the “right” formula. We compared 7 GAMs using R^2^ as model performance criterion (comparisons based on AICc led to the same interpretations, with the best model based on R^2^ obtaining an AICc weight >0.99). In all models we use the number of new patches populated between generations 100 and 120 as our “expansion velocity” response variable, as it was both after evolutionary changes were mostly complete and after velocity had seemingly reached an equilibrium (**Supplementary Material S4**). Since our landscape is made of discrete patches and the maximal possible expansion speed is one patch/generation, we analysed these data using binomial GAMs. There were some indications of underdispersion, likely because the probability of the expansion moving forward was not independent from one generation to the next, due to temporal autocorrelation in population sizes. More complex models accounting for this underdispersion gave similar conclusions (see archived code in **Data availability** for an example); we here only present the binomial GAMs results for simplicity. The simplest model, used as a null model of sorts, assumed expansion speed depended only on dispersal mortality (a categorical variable with three levels, **Table 1**) and not on any dispersal traits. The other models were divided equally into models that used trait information collected before the range expansion started, versus models that used trait information collected at generation 100, i.e. after genetic variation at the front was mostly spent (**Supplementary Material S4**). In both cases we analysed:

- (a) model assuming velocity depended on mortality, a spline effect of mean *d*_0_ at the range edge, and their interaction;
- (b) a model accounting for density-dependence by adding to (a) a spline effect of the mean difference *Δ*_*K*−0_ between *d*_0_ and *d*_*K*_ at the range front (and its interaction with mortality);
- (c) a model accounting for density-dependence by adding to (a) a spline effect of the mean difference *Δ*_*avg*−0_ between *d*_0_ and *d*_*avg*_ at the range front (and again its interaction with mortality).

### Genetic diversity

Another key aspect of pushed vs. pulled dynamics is the fate of genetic diversity, which is lost faster in pulled expansions compared to the “equivalent” pushed expansions (same *r*_0_ and *d*_0_) (Birzu et al., 2019). To better understand this, we analysed the effect of dispersal traits on time to genetic fixation, i.e. the first generation at which only one of the two neutral alleles is detected in the range front. Since we have grouped “survival” times, as we only collected information every 20 generations to save memory, we used cumulative ordinal regression models to analyse these data. In these models, the response variable is a categorical variable where the categories are ordered (here, along the time axis), and the category an observation belongs to can be seen as a coarse reflection of an unmeasured latent continuous response (here the actual time to fixation; Bürkner & Vuorre, 2019). When these models are fit with a complementary log-log link, they correspond to a proportional hazards model (e.g. Bürkner & Vuorre, 2019; Tutz & Schmid, 2016). We tested the same seven model structures as for velocity, although covariates were this time included “as is” and not as non-linear splines. Models were again compared using R^2^ as criteria. More accurately, we used a pseudo-R^2^ based on a linear regression between observed and mean predicted category (comparisons based on AICc led again to the same interpretations, with the best model based on R^2^ again obtaining an AICc weight >0.99). When analysing simulations with *r*_0_ = log(5), two simulations out of 2700 were excluded due to not having advanced (meaning they did not have any front to analyse). When analysing simulations with *r*_0_ = log(1.5), 64 replicate runs out of 2700 (so 2.4 %) were excluded, 21 because they did not advance, 43 because all populations went fully extinct before the end of the run.

### Trait evolution

Finally, to understand how trait evolution was influenced by initial conditions and dispersal costs, we focused on two traits: the density-dependence *Δ*_*avg*−0_ and the dispersal capacity *d*_*max*_. For *Δ*_*avg*−0_, we displayed the change in mean value in front patches after 100 generations as a fonction of the starting mean value for each combination of mortality, initial heritability, presence/absence of individual variation in dispersal traits at the start of the expansion, and used linear regressions to highlight the overall trends. For *d*_*max*_, we simply displayed the mean value at the front, since the initial mean value was the same (0.5) across all replicates.

## Results

### Results for *r*_0_ = log(5)

For both velocity and neutral genetic diversity, the best statistical models were the ones including information about both *d*_0_ and density-dependence (**Fig. 2**). For velocities, *R*^2^ was highest when we used traits after evolution to predict velocities, while for neutral diversity it was highest when we used traits measured before expansion (**Fig. 2**).

**Figure 2.**
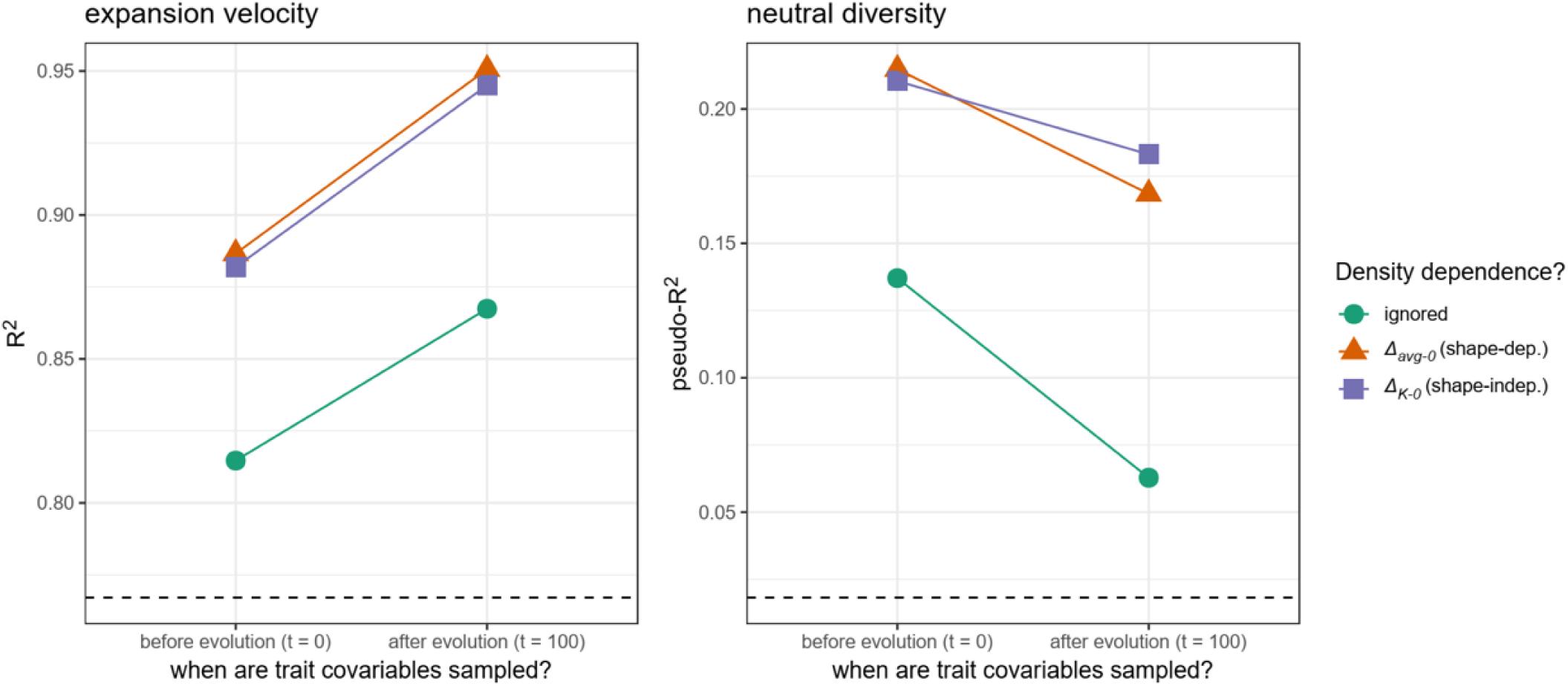
R^2^ of the different models used to analyse simulations outputs with respect to expansion velocity in the last 20 generations (left) and time to loss of neutral genetic diversity in the range front (right) (case where *r*_0_ = log(5), see **Supplementary Figures S5.1 and S5.2** for the case where *r*_0_ = log(1.5)). Dashed line: R^2^for the “baseline” model containing only dispersal mortality (all models include dispersal mortality, and its interactions with the other covariates when present).

Based on the best model, expansion velocity increased with *d*_0_ (**Fig. 3**), and for a given *d*_0_, expansions with positive density-dependent dispersal were consistently faster (**Fig. 3**). Increased mortality costs decreased velocity overall and reduced the effect of *d*_0_ and density-dependence on velocity, but did not change the overall direction of the effect.

**Figure 3.**
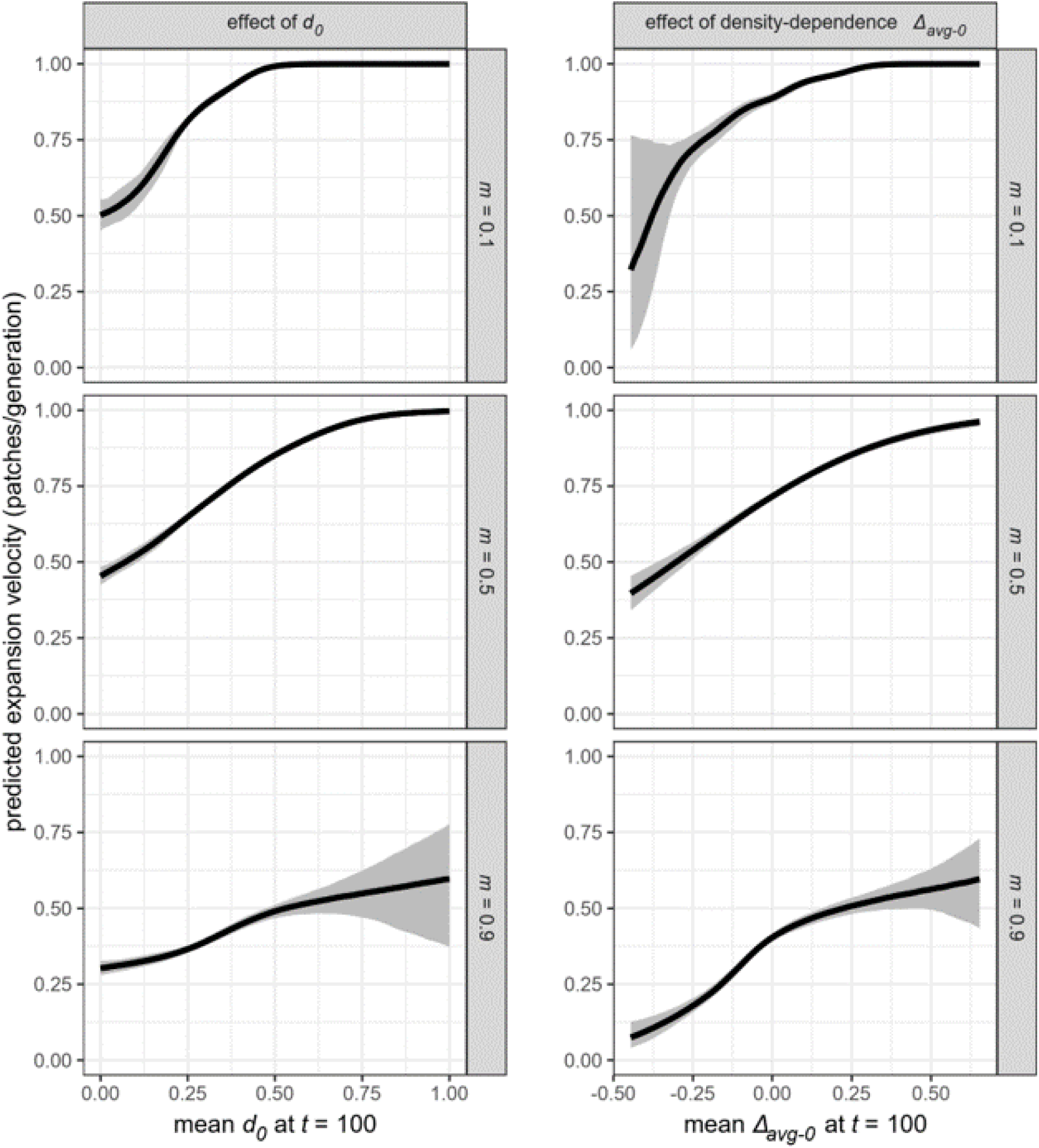
Predicted effect of *d*_0_ (left) and density-dependence *Δ*_*avg*−0_ (right) on expansion velocity, depending of dispersal mortality (case where *r*_0_ = log(5), see **Supplementary Figure S5.3** for the case where *r*_0_ = log(1.5)). Predictions are based on the best model **Fig. 2**; predictions for the effect of *d*_0_ are made assuming no density-dependence (*Δ*_*avg*−0_ =0), and predictions for the effect of *Δ*_*avg*−0_ are made setting *d*_0_ to its average value in the simulated dataset.

Higher *d*_0_ at the start of an expansion led to faster loss of neutral genetic diversity at the range edge, but only when mortality costs were high (**Fig. 4**). *d*_0_ being held equal, expansions that showed positive density-dependent dispersal maintained neutral diversity at the edge longer than those that did not, but only when dispersal costs were low or intermediate (**Fig. 4**). This effect of density-dependence was however reversed when dispersal costs were high (**Fig. 4**).

**Figure 4.**
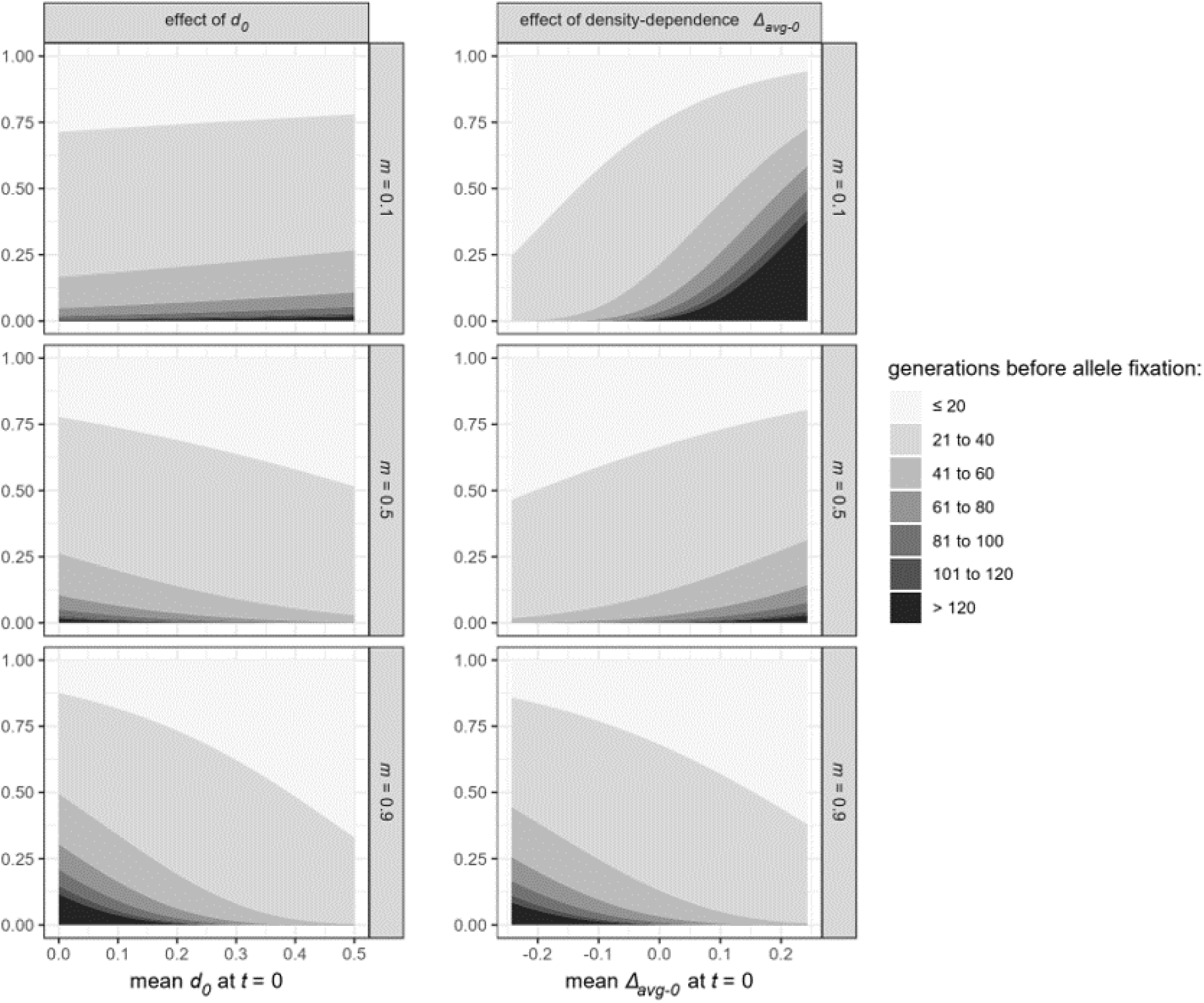
Predicted effect of *d*_0_ (left) and density-dependence *Δ*_*avg*−0_ (right) on the loss of neutral diversity at the range front, depending of dispersal mortality (case where *r*_0_ = log(5), see **Supplementary Figure S5.4** for the case where *r*_0_ = log(1.5)). Darker shades of grey indicate later fixation times. Predictions are based on the best model **Fig. 2**; predictions for the effect of *d*_0_ are made assuming no density-dependence (*Δ*_*avg*−0_ =0), and predictions for the effect of *Δ*_*avg*−0_ are made setting *d*_0_ to its average value in the simulated dataset.

When *d*_*max*_was allowed to evolve, it consistently evolved towards higher values at the range edge, whether or not other traits did evolve (**Fig. 5**). The strength of this response depended on heritability (evolutionary changes in *d*_*max*_were larger the more the trait was heritable) and mortality (changes were smaller the more dispersal was costly). Variance in outcomes was important and, although limited, cases of evolutionary decreases in *d*_*max*_could be observed (**Fig. 5**); they were more frequent with increased dispersal costs.

**Figure 5.**
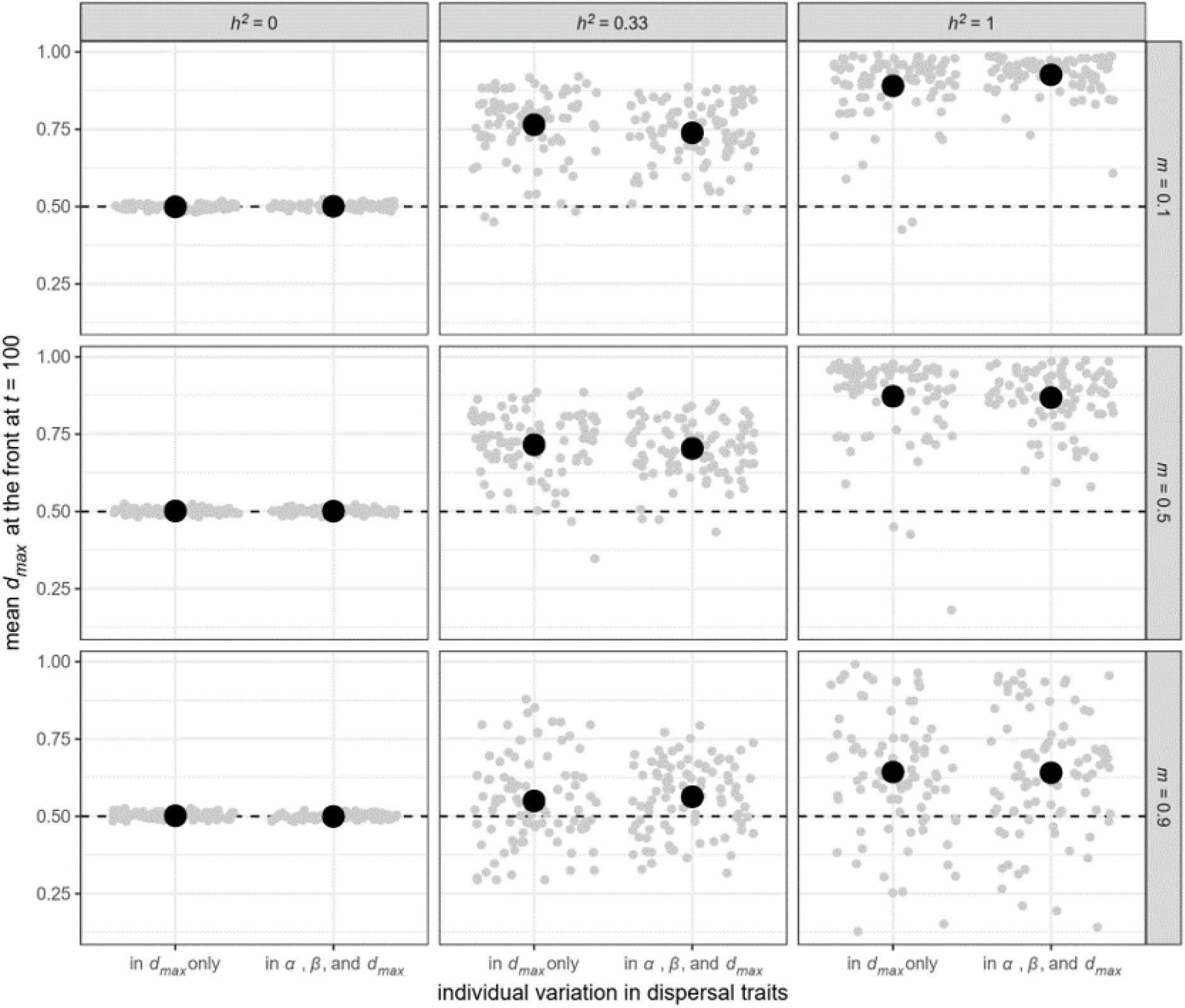
Evolutionary changes in dispersal capacity *d*_*max*_at the range front as a function of sources of phenotypic variation and dispersal costs (case where *r*_0_ = log(5), see **Supplementary Figure S5.5** for the case where *r*_0_ = log(1.5)). Grey dots are individual replicates, with the average values overlaid in black. Replicates with no individual variation in *d*_*max*_at *t* =0 are not displayed.

When only context-dependency (*α* and *β*) were allowed to evolve, changes in the density-dependence *Δ*_*avg*−0_ depended on heritability and dispersal costs, but overall leaned towards the evolution of more positive density-dependence, unless heritability was high and mortality costs were low (**Fig. 6A**). In that specific case, initially pushed expansions (positive density-dependence *Δ*_*avg*−0_) tended to lose that density-dependence with time (**Fig. 6A**, top right). When *only d*_*max*_was allowed to evolve, density-dependence tended to become stronger after evolution, with initially negative *Δ*_*avg*−0_ becoming more negative, and initially positive *Δ*_*avg*−0_ becoming more positive (**Fig. 6 B**). When both *d*_*max*_and context dependence were allowed to evolve, the general trend was to the evolution of positive density-dependence (**Fig. 6C**).

**Figure 6.**
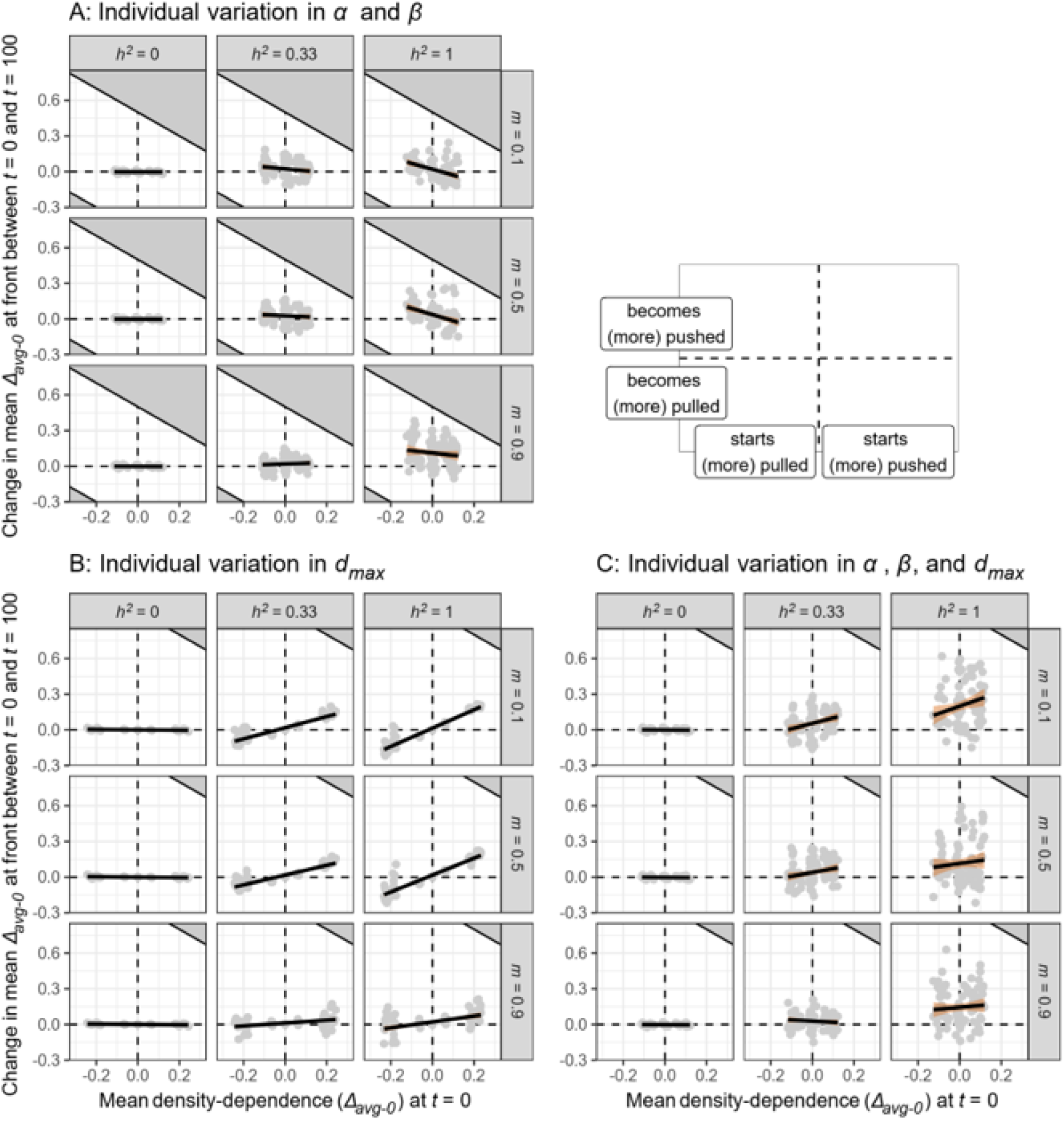
Evolutionary changes in density-dependence *Δ*_*avg*−0_ at the range front as a function of its initial value, sources of phenotypic variation and dispersal costs (case where *r*_0_ = log(5), see **Supplementary Figure S5.6** for the case where *r*_0_ = log(1.5)). Grey dots are individual replicates, with linear regression lines overlaid. Replicates with no individual variation in traits are not displayed. Grey-filled areas indicate impossible phenotypes, more specifically evolved values of *Δ*_*avg*−0_ that would imply dispersal rates higher than the maximal possible *d*_*max*_(1 if *d*_*max*_is variable among individuals, 0.5 if it is not).

For both *d*_*max*_and *Δ*_*avg*−0_, inter-replicate variation in evolutionary outcomes tended to increase the higher the initial trait heritability (**Figs 5, 6**).

### Sensitivity of results to growth rate

Most of the qualitative results described above were recovered when growth rate was set to log(1.5) (**Supplementary Material S5**) instead of log(5). However, when fecundity is lower, we find that:

- For genetic diversity, *R*^*2*^ and AICc diverge in which model structure is best (density-dependence included after evolution for the former, before evolution according to the latter; **Supplementary Figures S5.1 and S5.2**).
- When costs are high, high evolved values of *d*_0_ can reduce expansion speeds (**Supplementary Figure S5.3**).
- Dispersal capacity *d*_*max*_now evolves to lower values when costs are high enough (*m* =0.9) instead of consistently evolving to higher values (**Supplementary Figure S5.5**).
- As a result, expansions become less density-dependent when costs are high and only *d*_*max*_can evolve (**Supplementary Figure S5.6**)

We also note that when *r*_0_ = log(1.5), there remains substantially more genetic variation in traits after 100 generations than when *r*_0_ = log(5). However, the large majority of replicates still has lost most of its genetic variation (**Supplementary Material S4**).

## Discussion

A large number of experimental and observational studies have described how dispersal is often density-dependent, across a wide range of taxa (Harman et al., 2020; reviewed in Matthysen, 2005). In non-expanding metapopulations, many models have been developed to understand under what conditions density-dependent vs density-independent dispersal evolves and is maintained (e.g. Baines et al., 2020; Rodrigues & Johnstone, 2014; Travis et al., 1999 among many others). By contrast, until recently (Birzu et al., 2019), the implications of density-dependent dispersal for the dynamics of pushed vs. pulled expansions remained mostly unstudied, with research in that area focusing on Allee effects (e.g. Gandhi et al., 2016, 2019; Hallatschek & Nelson, 2008; Roques et al., 2012). In addition, although we know that dispersal traits (including matching traits shaping density-dependence) can evolve during range expansions (e.g. Chuang & Peterson, 2016; Fronhofer et al., 2017; Mishra et al., 2020), we do not know how sources of dispersal variation and dispersal evolution shape whether an expansion stays or become pushed or pulled. We here used simulation models as a first attempt to address this gap.

### Information on traits post-evolution is needed to (more) accurately predict pushed expansion speed but not neutral diversity

We find that our simulations generally behave as expected from pushed expansion theory (see e.g. Birzu et al., 2018, 2019), with some nuances however that may be key for its applicability in more ecologically realistic contexts. Importantly, we found that expansion speed was best predicted by the mean *d*_0_ and the average density-dependence of dispersal *after evolution* (**Fig. 2**). The more dispersal increased with density, the more expansions advanced faster than predicted from *d*_0_ alone and were therefore pushed (**Fig. 3**), as expected from theory (Birzu et al., 2019). Predictions based on *d*_0_ only, or predictions ignoring evolution and using trait values collected before the expansions performed consistently worse (**Fig. 2**). This is to some extent unsurprising, as any equilibrium speed would only be reached after dispersal evolution has occurred. However, this has practical implications: attempts to forecast expansions based on traits collected in non-expanding regions of the range only are likely to mispredict velocities whenever dispersal evolves at the edge (which may be often, Chuang & Peterson, 2016), unless one is able to also predict dispersal evolutionary dynamics themselves (for this, see below). Mortality costs strongly influenced the effect of dispersal traits on expansion speed (**Fig. 3**). While it did not change the direction of effect, information about environmental context (in particular connectivity) may be needed to make accurate predictions of the consequences of density-dependence on expansion speed.

As predicted by theory (e.g. Birzu et al., 2019), neutral genetic diversity loss was slowed down when dispersal was positively density-dependent (and therefore when expansions were pushed based on velocity) (**Fig. 4**). However, this was only true when dispersal costs were low or moderate. At high costs of dispersal, neutral diversity was actually lost slightly faster when dispersal increased with density (**Fig. 4**). We recover here a key result we highlighted in previous simulations, albeit without dispersal evolution (Dahirel, Bertin, Haond, et al., 2021): in some contexts, the responses of genetic diversity and expansion velocity to density-dependence are decoupled, and classifying an expansion as “pushed” based on one metric may not necessarily mean it will be “pushed” based on the other. This contrasts in a way with previous theoretical works implying that these two characteristics of pushed expansions are intrinsically linked (Birzu et al., 2018). Increased costs of dispersal are known to lead to lower genetic diversity during expansions (e.g. Mona et al., 2014). While we can expect that an increase in the number of dispersers/size of founding propagules may compensate for this, the net effect may still be negative if dispersal costs are too high (see e.g. Mona et al., 2014, for an example involving long-distance dispersal rather than density-dependent dispersal). Again, pushed expansion models that do not account for the diverse costs of dispersal (Bonte et al., 2012) in some way may then overgeneralize. This would lead to predictions that do not apply as effectively in contexts where costs of dispersal become important, for instance when species expand in highly fragmented environments.

Interestingly and contrary to expansion velocity, the best models explaining neutral genetic diversity loss used trait data collected at the start of the expansion, i.e. before trait evolution (**Fig. 2**). We believe this mostly reflects specific implementation details of our individual-based model, leading to generally fast loss of neutral diversity so starting dispersal traits had a disproportionate influence. Indeed, when fecundity was set to be lower, this discrepancy between velocity and diversity vanished, with both better explained by traits post-evolution (**Supplementary Figures S5.1, S5.2**). Further work is needed to see how reproduction and the structure of genetic diversity (ploidy, number of loci considered, number of alleles per locus), by influencing the baseline speed of diversity decay, can influence when dispersal traits have the strongest effect on diversity.

Finally, we note that while accounting for the shape of the density-dispersal function often leads to better predictions of the effect of density-dependence, models where the density-dependence was estimated based only on dispersal at *N* =0 and *N* = *K* still performed almost as well and sometimes better (**Fig. 2, Supplementary Figures S5.1, S5.2**). If confirmed, this is very interesting for future empirical studies. Indeed, it implies than most of the predictive benefits accrued by accounting for density-dependence can be obtained from a relatively small amount of dispersal data, even for pushed expansions.

### Sources of variation and costs of dispersal determine when dispersal evolution leads pushed expansions to become more pulled

Our simulations show that the strength and direction of the density-dependence seen at the range edge after evolution is to some extent predictable, if we have information about environmental characteristics (dispersal costs), initial trait values and sources of within-individual variation (**Fig. 6**). There still remains substantial amounts of unexplained variation (**Fig. 6**), especially at higher levels of initial heritability. This is in line with previous work showing that evolutionary stochasticity and serial founder events can lead to high variability in evolutionary outcomes and spread dynamics between initially identical or near-identical expansions (Dallas et al., 2020; Phillips, 2015; Weiss-Lehman et al., 2017). As already discussed by these authors and others, this puts an intrinsic limit on our ability to predict the evolutionary trajectories and spread of *individual* range expansions before they start.

Generally speaking, we find that Erm and Phillips (2020)’s key prediction, that pushed expansions tend to become pulled with time, is only fulfilled under a relatively narrow set of conditions when pushed dynamics are caused by density-dependent dispersal, rather than Allee effects as in their study. Simulated expansions starting from pushed dynamics (positive dispersal-density mean slope) only moved consistently towards more pulled dynamics when heritability was high, mortality was low and there was no individual variation in dispersal capacity i.e. *d*_*max*_(**Fig. 6**). When this conditions were not fulfilled, pushed expansions tended to stay pushed, and pulled expansions either stayed pulled or actually became pushed with time (**Fig. 6**). This is slightly surprising as, as detailed in the Introduction, Travis et al (2009)’s simulation study using the same dispersal function as ours concluded that dispersal become overall unconditionally high when it evolves at range edges. Our results show that the direct translation of that prediction into a prediction about pushed expansions, while intuitively appealing, must be done with extreme care, if at all. We do confirm that dispersal become overall less conditional at the edge (**Supplementary Material S6**). However, this must not be seen as synonymous with “expansions become less pushed”, as dispersal functions may be flat/unconditional over a large part of the range of densities yet still show positive density-dependence when compared to *d*_0_ (see e.g. **Fig. 1D**).

We believe the divergence between Erm and Phillips (2020) and our simulations hinges on the fact that the dispersal function is shaped by two types of parameters which can have conflicting effects on the absolute density-dependence reached after evolution:

- When only dispersal capacity i.e. *d*_*max*_can evolve, we find that pulled expansions become more pulled, and pushed expansions more pushed with time when *d*_*max*_increases due to spatial sorting (**Fig. 6B**), and the reverse in the few cases where *d*_*max*_decreases at the range edge (when high costs cannot be compensated by low fecundity, **Supplementary Figures S5.5 and S5.6**). As evolution leads to change in dispersal capacity along the range edge (**Fig. 5**), the *absolute* response to density can change even if the *relative* response does not (**Fig. 6**). Our results here contradict hypotheses put forward by Birzu et al. (2019), that evolutionary changes in dispersal motility would not influence “pushiness” since they would affect velocities the same way across the range of densities. We show here that this is not necessarily true. Increasing dispersal capacity increases the range of possible dispersal rates (Kun & Scheuring, 2006) (**Fig. 1**), and as a result increases the maximal potential amplitude of the density-dependence.
- When only dispersal matching traits (*α* and *β*) can evolve but not dispersal capacity, we find that initially pushed expansions are more likely to stay pushed when dispersal is more costly (**Fig. 6A**). This is fully in line with some aspects of existing dispersal theory, which predicts that higher costs of dispersal favour positive density-dependent dispersal strategies (Rodrigues & Johnstone, 2014; Travis et al., 1999). This also reproduces qualitatively results from a previous study involving two of the authors (Dahirel, Bertin, Haond, et al., 2021; Dahirel, Bertin, Calcagno, et al., 2021). The microwasp *Trichogramma brassicae* started range expansions with positive density-dependent dispersal and would thus demonstrate pushed dynamics in the absence of evolution. However, individuals at the edge rapidly evolved to develop more negative density-dependence, but only when connectivity was high (and thus presumably costs were low). Also, contrary to our initial hypotheses, we found that under some other contexts, evolution can lead to more, not less, positive density dependence in dispersal at range edges (**Fig. 6A**). That idea is also supported experimentally by at least one study (Mishra et al., 2020).

As a result, when both types of traits can evolve at the same time, their effects combine to shape the overall response (**Fig. 6**). The net effect will, in reality, likely depend on each trait’s heritability, trait variability and environmental context. However, it may be that, contrary to Erm and Phillips (2020) predictions for Allee effect-induced pushed expansions, pushed expansions caused by density-dependent dispersal are “resistant” to evolutionary change at the range edge in many circumstances. To be able to forecast if a given expansion will likely stay pushed, with the associated implications in terms of velocity or genetic diversity, we will therefore need better information about the degree of genetic variation in not only dispersal capacity, but also dispersal reaction norms (Saastamoinen et al., 2018), information that is currently lacking for many species, including expanding ones.

### Conclusion: towards even more realistic representations of phenotypic variation

Our simulation study shows that whether an expansion stays or become pushed during spatial evolution is to some extent predictable from initial conditions, but information about dispersal costs (and probably more generally environmental conditions) is needed to make correct predictions. Predictions also depend on whether individuals vary in dispersal capacity, relative response to density, or both. While our model is arguably more complex than some of the existing ones, especially due to the addition of evolution, it still makes some key simplifications, which may need to be reexamined to determine whether they influence our results.

First, we used asexual haploid virtual organisms and each dispersal trait was shaped by one locus only; evolutionary dynamics might be influenced by the reproductive system, or the number of loci involved in inheritance and their genetic architecture (Saastamoinen et al., 2018). Recent simulations suggest that while these may influence the strength of evolutionary change during expansion (and its effect on expansion velocity), they might not change the overall direction of the evolutionary response (Weiss-Lehman & Shaw, 2021). These results remain however to be confirmed in the context of pushed expansions and density-dependent dispersal. We also ignored mutation for simplicity and focused only on the fate of initial standing variation. Work by Erm and Phillips (2020) suggests that adding mutation would merely accelerate/accentuate trait differentiation compared to what we observed, as it would provide variation allowing spatial sorting to continue.

Second and most critically, we only studied the effect of dispersal evolution. Whether or not an expansion is pushed, and to what degree, depends on both dispersal and population growth (Birzu et al., 2019), and the latter can be further subdivided in its fundamental life history components, fecundity and survival. However, dispersal, fecundity and survival can all vary among individuals, and can all be density-dependent at the same time, potentially in ways that could cancel each other out. They may also be correlated in syndromes at the within- and/or among-species level (Beckman et al., 2018; Guerra, 2011; Jacob et al., 2019; Ochocki et al., 2020; Ronce & Clobert, 2012), and these syndromes may shape and constrain the evolution of their constituent traits (Ronce & Clobert, 2012; Wright et al., 2019), including during range expansions (Ochocki et al., 2020; Urquhart & Williams, 2021). In addition, theoretical work even suggests that density fluctuations themselves are one of the root evolutionary drivers shaping the (co)variation of life history traits, and potentially their association with behaviours such as dispersal (Wright et al., 2019, 2020). How the complexities of phenotypic structure and trait coevolution drive the net effect of population density on spread, influencing the dynamics of pushed vs pulled expansions and our ability to predict them, remain to be discovered.

## Data availability

Netlogo model code and R scripts to reproduce all analyses presented in this manuscript are available on Github (https://github.com/mdahirel/pushed-pulled-2020-heritability-IBM) and archived in Zenodo (https://doi.org/10.5281/zenodo.5830993). Simulation outputs are also archived in Zenodo (https://doi.org/10.5281/zenodo.5830996).

## Author contributions

Funding: EV; initial manuscript idea: MD and EV; simulation model development: MD and CG; simulated data analysis: MD; initial manuscript draft: MD. All authors read and edited the manuscript, and approved the final version.

