## Supplementary Material for "Individual variation in dispersal, and its sources, shape the fate of pushed vs. pulled range expansions"

#### S1 - Detailed description of the individual-based model in the ODD format

##### Purpose and patterns

The model simulates asexual individuals reproducing and dispersing in a linear landscape following an initial introduction, in order to approximate processes at play during natural and experimental range expansions. The aim is to gather insights into how evolutionary dynamics (neutral diversity and especially dispersal-related traits) are shaped by trait distribution (means and variances) at the start of the expansion, and how they influence expansion velocity in turn. We specifically focus on traits driving the position of expansions on the pushed/pulled continuum (Birzu et al., 2018, 2019), namely traits describing the shape of the density-dispersal function. Importantly, dispersal and reproduction are stochastic. The focus of the model is on studying the effects of initial trait variation at introduction; as such there is no mutation, for simplicity.

##### Entities, state variables and scales

Individuals are haploid and reproduce asexually, and can vary in:

- their location in the landscape ($x$ coordinate only, since we simulate linear landscapes. $x$ can only take $\geq 0$ integer values);
- their dispersal probability $d$ (real between 0 and 1) and the underlying dispersal traits ($d_{max}$, $\alpha$, $\beta$). Each trait is the (transformed) sum of an additive genetic component ($\text{logit}(d_{max})_{[a]}$, $\alpha_{[a]}$, $\beta_{[a]}$) and of a noise/residual part ($\text{logit}(d_{max})_{[r]}$, $\alpha_{[r]}$, $\beta_{[r]}$). All these components are reals. See **Initialization** and **Submodels** below for details about the dispersal function;
- the allelic value at a neutral locus $\gamma$ (binary, 0 or 1);
- their life stage (juvenile, adult, coded as a binary variable);
- their fecundity $F$ ($\geq 0$ integer).

We note that the model as it is coded and made available (**Data availability**) is more general and allows the use of diploid, sexually reproducing individuals, with some slight adaptations in the reproduction submodel. For simplicity we only use and describe the "haploid asexual" scenario here.

Individuals live in discrete patches, which form the spatial units of our model. The model world is one-dimensional, and has closed boundaries. Patches can be described by:

- their location in the landscape ($x$ coordinate; $\geq 0$ integer values);

- their population size at the last count $N$ ($\geq 0$ integer; population counts occur at points of the cycle where only adults are present);
- their carrying capacity $K$ ($\geq 0$ integer; in the present study, $K$ is fixed and constant across all patches);
- a series of variables summarizing the trait distributions of the individuals born in the patch (see **Submodels**).

One time step in the model represents one generation of the simulated species' life cycle, and each individual grid cell corresponds to an individual patch. The potentially hostile matrix between patches is abstracted out (there are no "matrix" grid cells) and its effects are summarised in a global environmental variable, dispersal mortality $m$. The spatial and temporal extents of the simulations are not fixed and can be set up based on the needs of the simulation experiments.

#### Process overview and scheduling

Generations start at the point of the life cycle where only pre-dispersal, pre-reproduction adults are present. The schedule of a generation can be divided in 6 processes: *initial Population count*, *Summary statistics*, *Dispersal*, *second Population count*, *Reproduction*, and *Death*. All individuals and patches must go through one process before the model proceeds to the next. The order in which individuals/patches go through a given process is random.

- *Population count:* Individuals at each patch are counted to update the patch population sizes $N$ and the neutral allele frequencies for each patch.
- *Summary statistics:* Patches estimate a series of summary variables describing the trait distribution of individuals they currently harbour. This is done because exporting and analyzing individual-level traits would be computationally expensive. A detailed list of these summary variables is provided in **Submodels** below.
- *Dispersal:* Adult individuals may disperse with a probability $d$ depending on their individual traits ($d_{max}$, $\alpha$, $\beta$) and local population size $N$ **at the start of the dispersal phase**. The dispersal model is taken from Kun and Scheuring (2006) and described in **Submodels** below. If an individual attempts to disperse, it dies during the attempt with a probability $m$. In practice, we draw random values from a Uniform$(0,1)$ distribution; if the draw is lower than $d$ (respectively $m$), the individual disperses (respectively dies). Surviving dispersers are randomly moved to one of the nearest neighbouring patches; that is, the maximal dispersal distance is of 1 patch.
- *Second population count:* The population sizes $N$ and neutral alleles frequencies at each patch are updated again.
- *Reproduction:* Each remaining adult then produces $F$ juveniles, with $F$ drawn from a Poisson distribution and depending on the general growth rate $r_0$, $K$ and $N$. Juveniles are born in the patch currently occupied by their parent (same $x$), and also inherit their genetic values for dispersal-related and neutral loci. Their values for the noise part of the dispersal traits are redrawn at random. The exact fecundity model and inheritance procedures are described in **Submodels** below.
- *Death and end of cycle:* All adults die; juveniles then become adults.

The model then starts the next generation. We chose to make dispersal precede reproduction, so that our model simulates *natal dispersal*, i.e. the movement from birth site to the first reproduction site, arguably the most studied and best understood type of dispersal.

#### Design concepts

**Basic principles:** This model follows previous attempts to model evolution in range expansions using an individual-based simulation framework (e.g. Travis et al., 2009), including in the context of pushed expansions (e.g. Erm & Phillips, 2020). This model is a theoretical investigation tool and is not built to make any real-world quantitative prediction, but to develop new qualitative insights about evolutionary dynamics during range expansions. It differentiates itself from previous modelling studies of pushed expansions in an ecological/evolutionary context by:

- the use of a more flexible dispersal function (Kun & Scheuring, 2006).
- a focus on "small" population sizes that are realistic for many "macroscopic" organisms, giving a more important role to discreteness, drift and stochasticity in general, compared to models assuming much larger population sizes either implicitly (e.g. when using continuum models) or explicitly (Birzu et al., 2019).
- allowing for varying levels of trait heritability, where most models either ignore evolutionary dynamics in focal traits (e.g. Birzu et al., 2019) or assume all variation is genetic (e.g. Erm & Phillips, 2020).

**Emergence:** The number and distribution of individuals and alleles in space are the emergent result of individual dispersal and reproduction, which are stochastic and themselves depend, recursively, on the distribution of individuals in space. Most if not all variables of interest are summaries from these distributions.

**Adaptation:** As population size varies in space and time along the range expansion, dispersal is costly, and fitness is density-dependent (see **Submodels**), dispersal can evolve in an adaptive way in replicates where dispersal traits are heritable.

**Objectives:** There is no direct objective-seeking in this model, in the sense that individuals do not adjust their dispersal behaviour *for* increasing their expected fitness. However, the model allows for evolution; in these cases, natural selection based on differential reproductive success may lead lineages to maximize their reproductive success through time.

**Learning:** Individuals do not learn (see **Sensing**).

**Prediction:** There is no prediction component to individuals' choices: individuals behave only based on currently available information, including their phenotype.

**Sensing:** In addition to its own trait values, the only information an individual can sense and use is the (last updated) population size $N$ in the patch it is currently in. Individuals automatically and accurately know $N$ when needed, and immediately forget it afterwards (there is no memory).

**Interaction:** Individuals do not interact directly with each other. Individuals in the same patch may influence each other indirectly, through the patch population size variable $N$ which plays a role in both individual dispersal and reproduction.

**Stochasticity:** Dispersal and reproduction are both stochastic. First, for dispersal, the individual probability of dispersing depends on three traits, the values of which are determined stochastically. The genetic components are drawn from Normal distributions at the start of the experiments (and then transmitted deterministically), and the noise components are drawn from Normal distributions at the birth of every individual. Second, once the individual dispersal probability is set, whether or not an individual actually disperses, and whether or not it dies if it disperses, is in effect determined by Bernoulli trials (implemented by determining if a (pseudo-)random number between 0 and 1 is higher or lower than the relevant probability). Finally, the actual fecundity of each individual is a random draw from the relevant Poisson distribution. See **Initialization** and **Submodels** for more details.

**Collectives:** Individuals do not belong to any kind of identifiable social group or collective *per se*. Individuals in the same discrete patch nonetheless influence each other through $N$ (see **Interaction**)

**Observation:** For memory reasons, it is impractical to save the state of every individual. We instead measure and record patch-level metrics, which convey information about population size, neutral allele frequencies and the means and variances of variables summarizing the dispersal-density functions of individuals born in the patch (see **Submodels**). The user can choose to save these metrics every generation, or for only a subset of generations of interest.

#### Initialization

Each expansion starts by creating the landscape (which, as we use nearest-neighbour dispersal, can be any length higher than the number of generations planned for the run), and setting the carrying capacity of all patches to $K$. We then introduce $K$ adult individuals that have not yet dispersed or reproduced to the left-most patch of the landscape ($x = 0$). All other patches are empty. Individual traits and alleles are then initialized as follows:

- the value (0 or 1) at the neutral allele $\gamma$ is drawn from Bernoulli(0.5).
- for each dispersal trait $z$ (logit($d_{max}$), $\alpha$, $\beta$), the genetic component is drawn from Normal($\overline{z}_{t=0}$, $\sqrt{h^2 V_{P[z]}}$) , the noise component from Normal(0, $\sqrt{(1-h^2)V_{P[z]}}$) , where $\overline{z}_{t=0}$ is the initial mean trait value, $h^2$ is the initial trait heritability and $V_{P[z]}$ the initial phenotypic variance. The trait value is then obtained by summing the genetic and noise components, and in the case of $d_{max}$, back-transforming from the logit scale.
- we calculate secondary individual-level statistics from $d_{max}$, $\alpha$ and $\beta$ (see the Dispersal submodel description below).

#### Input data

The model is self-sufficient and does not use external input data to represent time-varying processes.

#### Submodels

**Dispersal:** during the dispersal phase, an individual leaves its natal patch with a probability $d$, which depends on individual traits and local population size $N$ as measured at the start of the dispersal phase (here expressed relative to the constant $K$) based on Kun and Scheuring (2006)'s model (see also Travis et al., 2009):

$d_N = \frac{d_{max}}{1+\exp(-\alpha(\frac{N}{K}-\beta))}$.

Compared to other dispersal models (such as those reviewed in Hovestadt et al., 2010), this three-parameter model has the advantage of flexibility, can model a wide variety of realistic dispersal curves, and allows for both negative and positive density-dependent dispersal. In addition, all three parameters have an ecologically relevant interpretation:

- $d_{max}$ is the maximal dispersal rate, and can be seen as a measure of dispersal capacity. Note that depending on the other parameters, $d_{max}$ may not be reachable at any of the population sizes actually experienced during the simulation.
- $\alpha$ is the slope of the dispersal-density relationship and describes the "strength" of the response to population density (relative to $d_{max}$). Whether $\alpha$ is positive or negative determines whether dispersal increases or decreases with population size. Depending on $\beta$, $\alpha$ may or may not influence meaningfully the actual dispersal strategy at the population sizes experienced during the simulation.

- $\beta$ is the inflection point of the dispersal function, i.e. the density at which $d = 0.5 \times d_{max}$. It can be interpreted as a response threshold, i.e. how high (or low) density must get for the individual to substantially alter its dispersal response. We allow $\beta$ to be negative; this can be interpreted as individuals being so sensitive to density that they already are close to their high density dispersal behaviour at low densities. For our purpose, this is the only practical way the model can produce density-independent functions where $d \approx d_{max}$ (as setting $\alpha$ to 0 actually sets $d$ to be constant, but $= 0.5 \times d_{max}$). Another way is theoretically possible: allowing $\beta$ to be close to 0 or $K$ but positive, and $|\alpha|$ to be so high (for a given $K$) that dispersal goes from 0 to $d_{max}$, or the opposite, between $N = 0$ and $N = 1$ (as in e.g. Travis et al., 2009). However, due to how $\alpha$ influences the shape of the dispersal function, Normal distributions of initial $\alpha$ that allow for such high $|\alpha|$ would be strongly biased against shallower slopes (see **Supplementary Material S2**).

Following the terminology in Cote et al (2017), variation in $d_{max}$ can be interpreted as encoding variation in dispersal enhancing or enabling traits, while variation in $\alpha$ or $\beta$ corresponds to variation in dispersal matching traits.

Once we know the $d_{max}$, $\alpha$ and $\beta$ of an individual, we additionally calculate five values at the individual-level:

- $d_0$, the hypothetical dispersal rate at $N = 0$. The theoretical distinction between pulled and pushed expansions hinges on whether expansion move as fast as expected from $d_0$ or faster (e.g. Birzu et al., 2019).
- $d_K$, the expected dispersal rate at $N = K$.
- $d_{avg}$, the average dispersal over the range of densities $0 - K$. For computational reasons, we approximate it by calculating $d_N$ every $0.1K$ from 0 to $K$.
- Two measures of the strength of density-dependence $\Delta_{K-0} = d_K - d_0$ and $\Delta_{avg-0} = d_{avg} - d_0$. The former is independent of the shape of the dispersal function between 0 and $K$, while the latter accounts for it at least in part.

**Reproduction:** Each adult individual still living during the reproduction stage produces $F$ offspring, with $F \sim \text{Poisson}(\lambda)$ and the average offspring number $\lambda$ depending on local population size $N$ based on a Ricker model:

$\lambda_N = \mathrm{e}^{r_N}, r_N = r_0(1 - N/K)$,

where $r_0$ is the hypothetical *per capita* growth rate at $N = 0$.

Parent transmit their genetic values (dispersal-related: $\text{logit}(d_{max})_{[a]}$, $\alpha_{[a]}$, $\beta_{[a]}$ and neutral: $\gamma$) directly to their offspring without any mutation. Offspring then draw random values of noise components $\text{logit}(d_{max})_{[r]}$, $\alpha_{[r]}$, $\beta_{[r]}$ from $\text{Normal}(0, \sqrt{V_{R[z]}})$, where $z$ is the trait in question. For each trait $z$, the "random noise" variance $V_R$ is equal to $(1-h^2)V_{P[z]}$ , where $h^2$ is the *initial* heritability at $t = 0$ and $V_P$ the *initial* phenotypic variance.

**Observation:**

The code estimates and reports a variety of summary variables at the patch level, and can easily be modified to add more. Of potential interest are the following:

- the means of $d_{max}$, $d_0$, $d_{avg}$, $d_K$, $\Delta_{K-0}$, $\Delta_{avg-0}$ for all individuals born in a patch.
- the variances of the genetic components $\text{logit}(d_{max})_{[a]}$, $\alpha_{[a]}$, $\beta_{[a]}$ for all individuals born in a patch.
- population sizes $N$ pre- and post-dispersal phase.
- the frequencies of the two neutral alleles in a patch, both pre- and post-dispersal phase.

#### S2 - Effect of the magnitude of initial variation in $\alpha$ on density dependence

In the ODD description above, we briefly discuss how allowing for negative $\beta$ is the only way we have to create mostly flat dispersal-density function where dispersal is close to $d_{max}$. We briefly explain that one seeming alternative, allowing very large $|\alpha|$ values so that dispersal goes from 0 to $d_{max}$ in less than 1 individual, is not viable. We expand this argument here.

Using very large $\alpha$ means we'd need the initial trait distribution to contain both these very large $\alpha$ (positive and/or negative) *and* $\alpha$ close to 0, for evolution to be possible (in the absence of mutation at least). It's easy to demonstrate that distributions that fulfill this requirement are actually biased towards very sharp density-dispersal functions. If we assume $K = 250$ as in our simulations, then $\alpha = 1000$ (with $\beta$ very close to 0) is needed to generate one of these nearly flat functions:

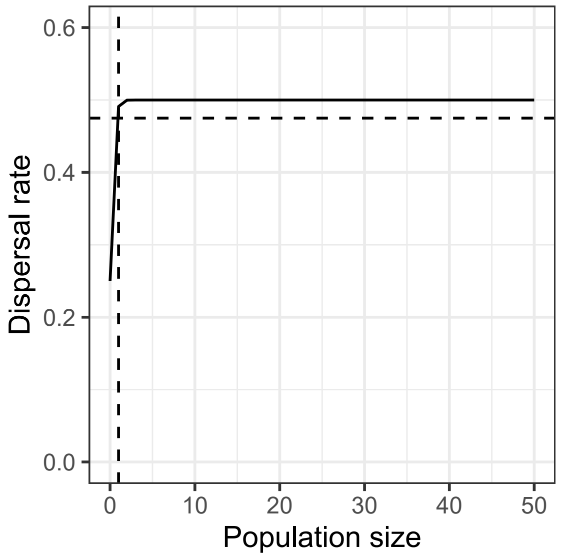


**Figure S2.1**. Example of how one can produce a functionally flat density-dispersal function using a very sharp slope, but very close to $N = 0$. $d_{max} = 0.5$, $\alpha = 1000$, $\beta = 10^{-15}$. The dashed lines mark $N = 1$ (vertical) and $d_N = 0.95 \times d_{max}$ (horizontal).

So, even after setting $\beta$ absurdly close to 0, we still need $\alpha$ in the neighbourhood of 1000 to be able to produce such shapes (when, as a reminder, $\alpha = 6$ is needed to get a slope that takes a population increase of $K$ individuals to go from 5% to 95% of $d_{max}$, so a reasonably shallow, but not too shallow function). An initial distribution of $\alpha$ of $\text{Normal}(0, 500)$ would hit these extreme $\alpha$ with a low but non-negligible probability (about 5%). And we know (see main text) that $6/\alpha$ approximately expresses (in units of $K$) how sharp the increase in dispersal with density is. Let's have a look at the distribution of $6/|\alpha|$ implied by $\alpha \sim \text{Normal}(0, 500)$ :

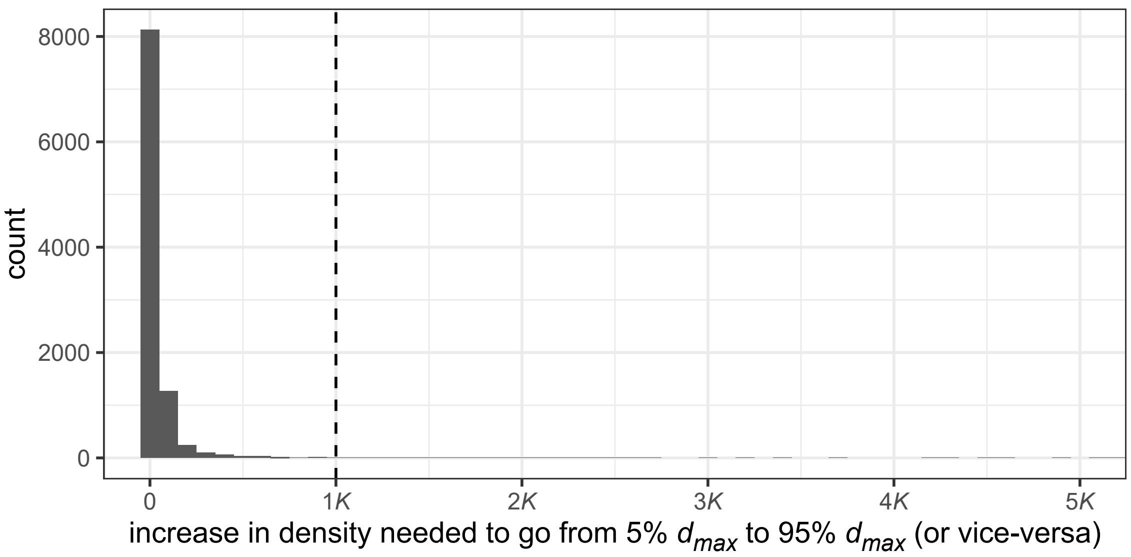


**Figure S2.2**. Distribution of the increases in density needed to go from 5% of $d_{max}$ to 95%, assuming $\alpha \sim \text{Normal}(0, 500)$. Histogram drawn based on 10000 samples.

We can see that this initial distribution of $\alpha$ would mostly only allow sharp slopes, that take less than 20-50%$K$ to go from (nearly) 0 to (nearly) $d_{max}$, and shallower slopes would only be present at much lower frequencies. This is not a choice that is ecologically sound in our opinion, hence why we rejected "extreme $\alpha$" as a method to generate $d_N \approx d_{max}$ flat functions.

### S3 - How do we define the "range front?"

We are interested in how traits at the front of the expansion are shaped by, and shape, the expansion process. To do that right, we need an operational definition of what is the front. A "traditional" one is the range of patches where a density gradient is seen, i.e. where population density has not yet reached its equilibrium (e.g. Gandhi et al., 2016; Lewis et al., 2016). Let's have a look:

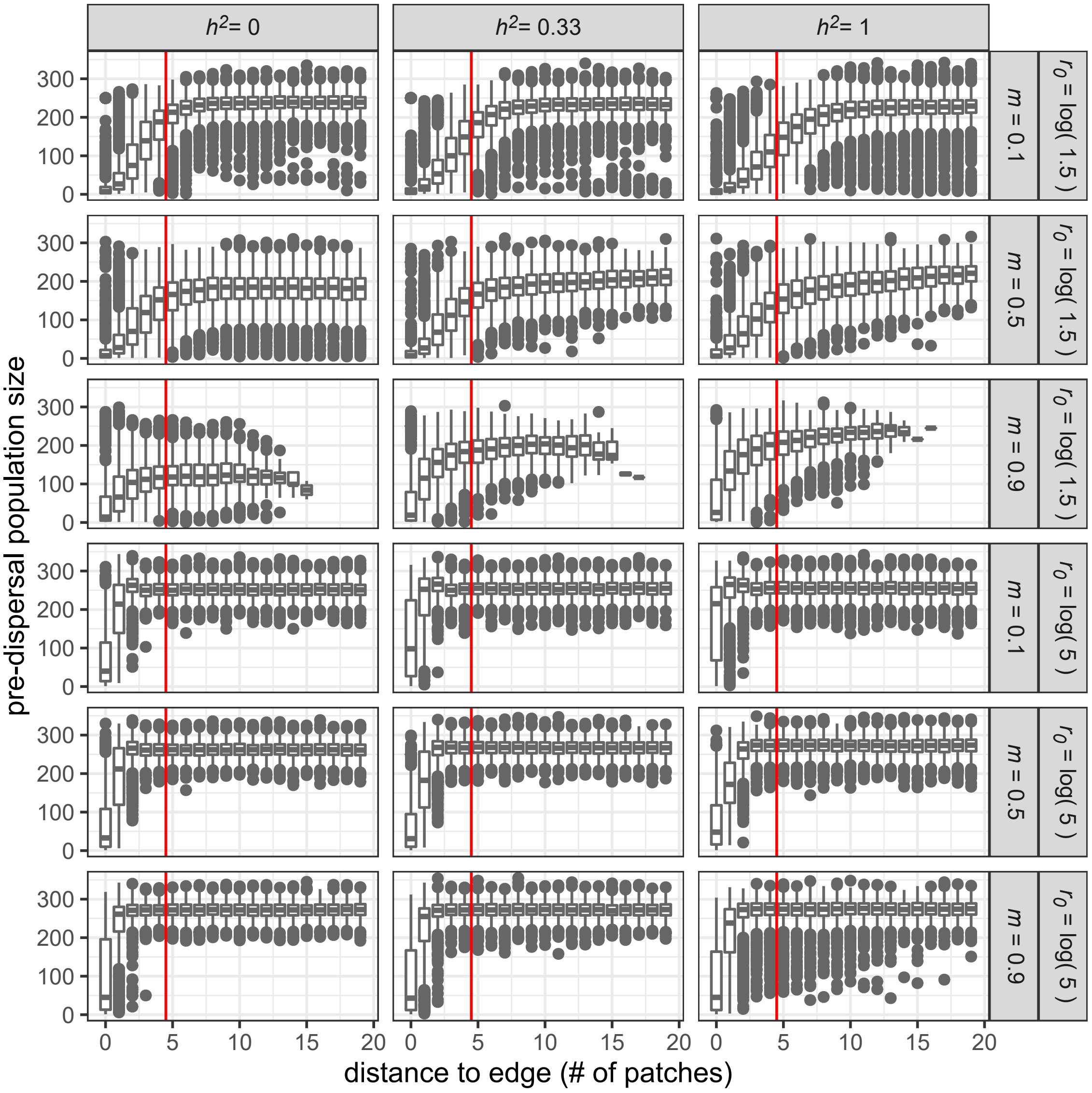


**Figure S3.1**. Predispersal population size as a function of distance to the range edge (only populations up to 20 patches from the edge at the time of sampling included). The vertical line separate populations < 5 patches from the edge from the others.

We can see that a threshold where we consider all populations < 5 patches from the edge as belonging to the range front is a good compromise that works across conditions. Narrower fronts would probably work for the high fecundity scenarios, but would not work for the low fecundity ones. In addition, a narrower threshold would limit the number of individuals included, reducing the precision of our trait means, while a wider one would lead us to fronts that are dominated by what are clearly non-front patches when fecundity is high.

### S4 - Checking velocity and evolutionary convergence

#### Checking velocity convergence

If an expansion has converged to its asymptotic speed after a time $t$, then it means that its speed over the full duration of the simulation (as long as it is $>t$) and its speed over any time interval *after t* should be identical (± stochastic noise, of course).

We find that convergence in expansion velocities is fast, and speeds measured at or after 100 generations can reasonably used as measures of long-run speed:

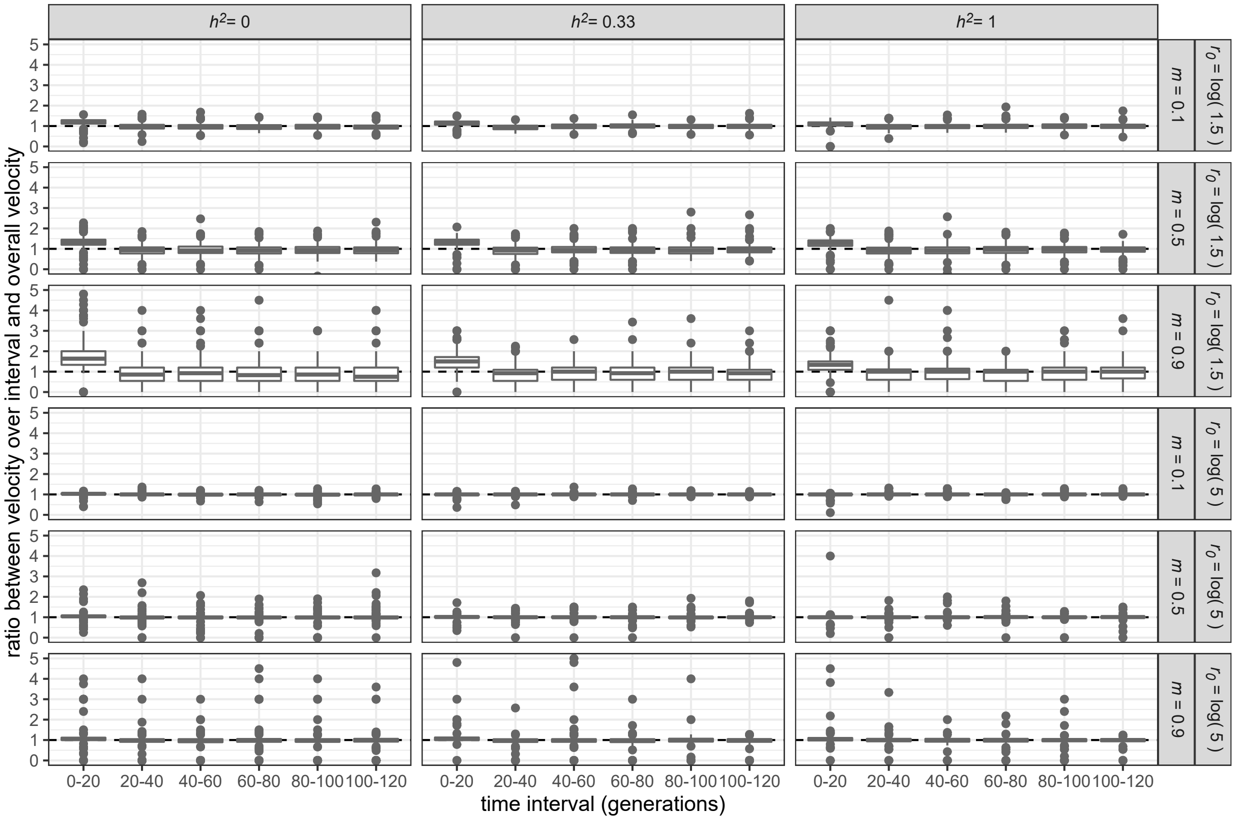


**Figure S4.1**. Convergence of expansion velocity towards an equilibrium. Ratio of the velocity observed over a 20-generation interval and the velocity over the entire 120-generation run. y-axis is cut to [0;5] to better showcase the interquartile range.

#### Checking evolutionary convergence

We can then do a similar check for evolutionary stability at the edge. Because there is always some genetic variation remaining, let's use "only 5% of the initial genetic variation in traits remaining on average in front patches" as an arbitrary cutoff:

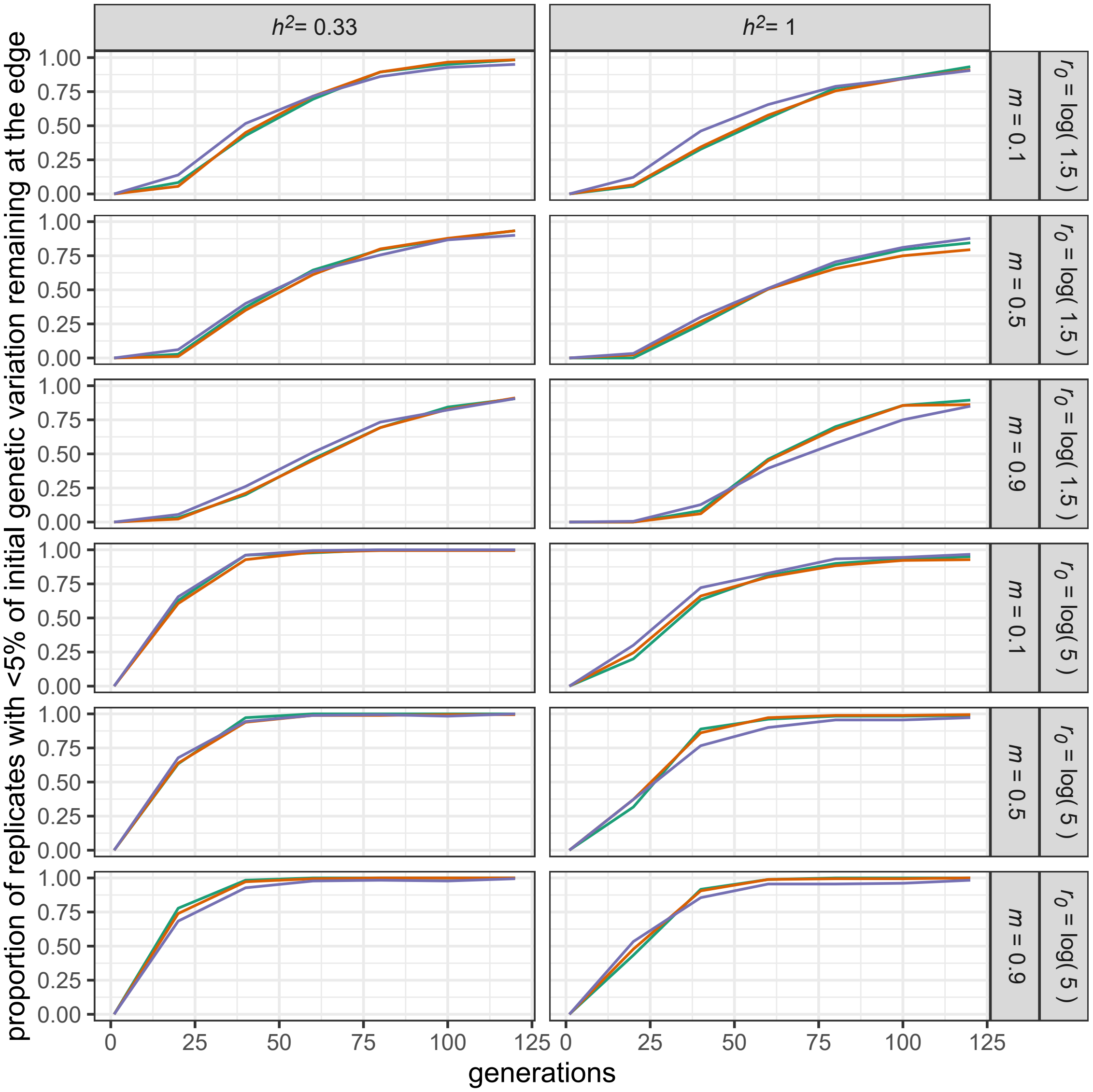


**Figure S4.2**. Proportion of replicates that have "spent" their initial genetic variation, as a function of time. Within a panel, each line represents a different trait, with midpoint ($\beta$) in green, slope ($\alpha$) in orange and $d_{max}$ in blue (the trajectories for all three traits are nearly identical).

Using our operational definition of evolutionary "stability," we note that a very large majority of replicates have converged by the end of the experiment, especially in high fecundity replicates. In addition, we note that our definition of evolutionary stability excludes *de facto* cases of divergent selection, so the % of "evolutionary stable" expansions by $t$=100 generations, say, is possibly even higher.

### S5 - Results for fecundity = 1.5

Here we reproduce the analyses and figures displayed in the main text Results, but for the low-fecundity scenarios. Note that 65 out of 2700 replicates were excluded when fitting the velocity models, due to going fully extinct before the window used for velocity models (between $t = 100$ and $t = 120$) or because they had negative speeds during that window. We chose to exclude a very small percentage and stay consistent to the analyses in the main text, rather than change the analysis method for the Supplementary analyses.

Regarding model selection, it is interesting to see here that, while the conclusions are the same as for fecundity = 5 for velocity (the best model includes density-dependence post-evolution, whether we use $R^2$ or AICc as a criterion), things are a little bit more complex for genetic diversity. Here, while both evaluation metrics agree on the fact the best model must include density-dependence, they disagree on whether it should be post- or pre- evolution. Note that we still decided to build **Fig. S5.4** using traits measured pre-evolution, to be comparable with the main text **Fig. 4**.

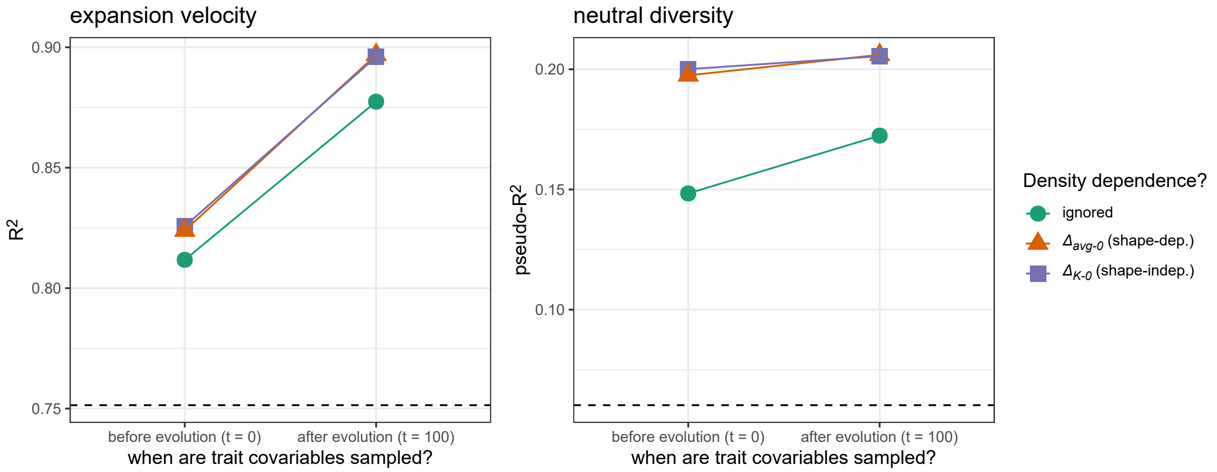


**Figure S5.1**. $R^2$ of the different models used to analyse simulations outputs with respect to expansion velocity in the last 20 generations (left) and time to loss of neutral genetic diversity in the range front (right) (case where $r_0 = \log(1.5)$, see **main text Fig. 2** for the case where $r_0 = \log(5)$). Dashed line: $R^2$for the "baseline" model containing only dispersal mortality (all models include dispersal mortality, and its interactions with the other covariates when present).

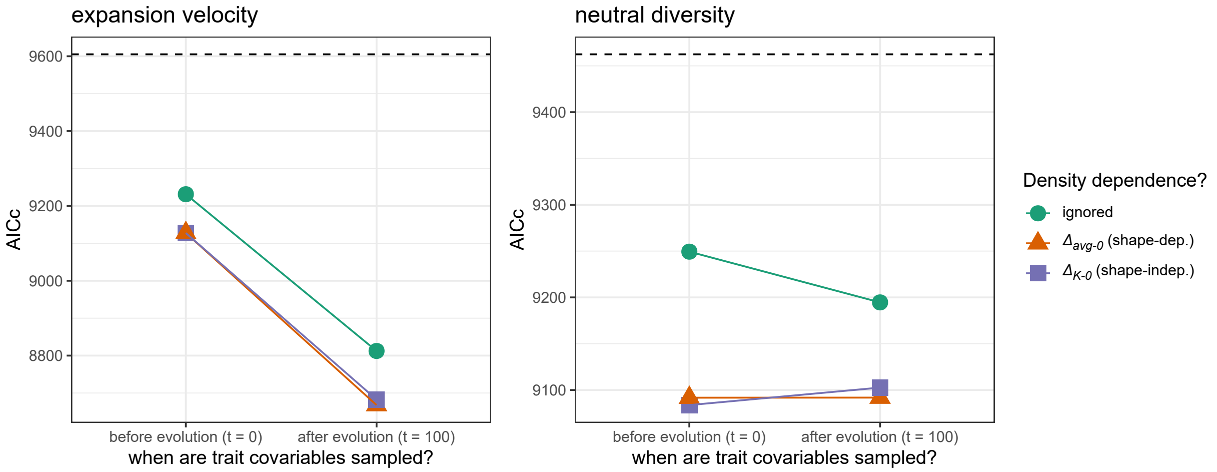


**Figure S5.2**. Same as **Figure S5.1** except with AICc as the model evaluation criterion. Note that this time, *lower* values denote better models.

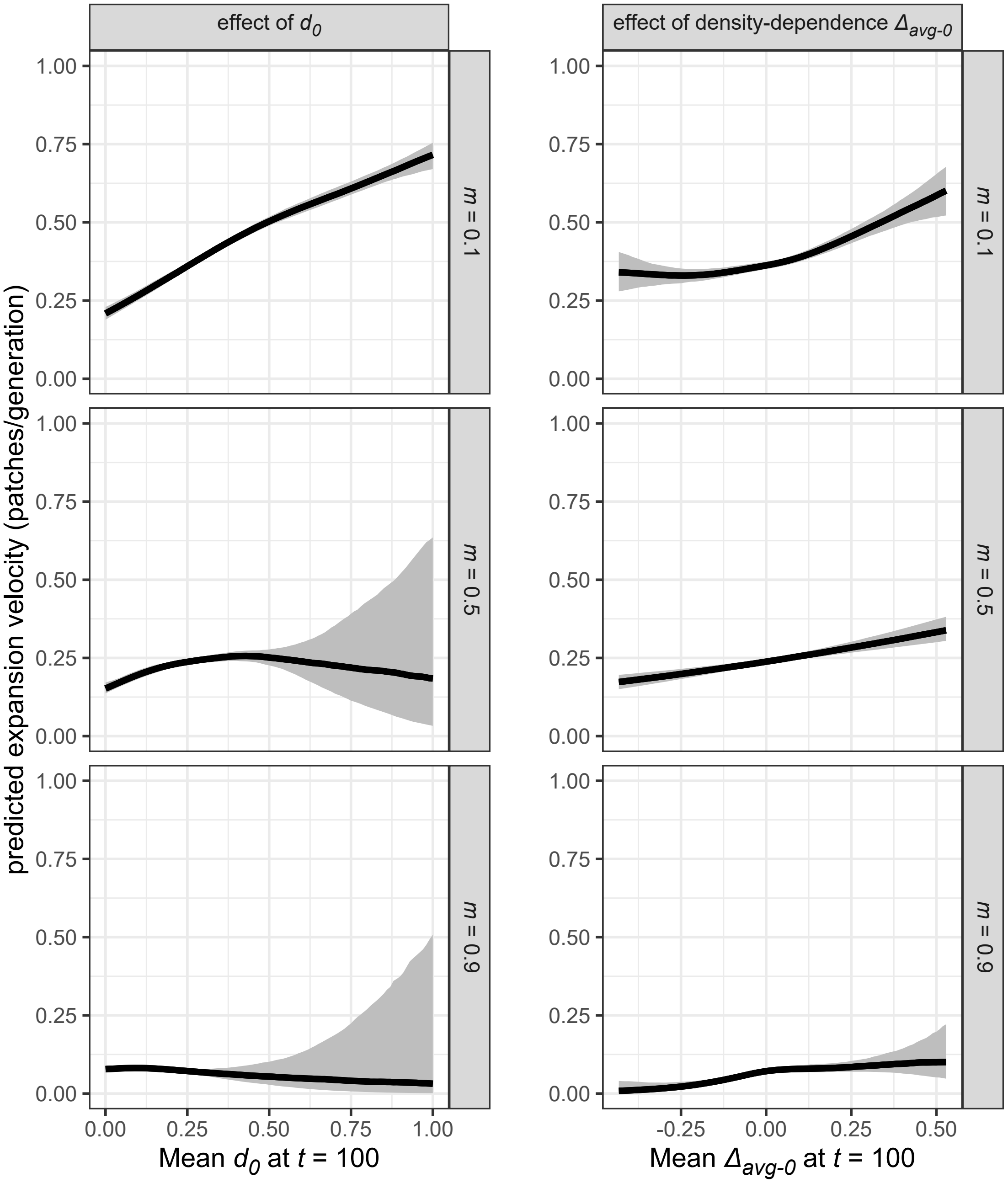


**Figure S5.3**. Predicted effect of $d_0$ (left) and density-dependence $\Delta_{avg-0}$ (right) on expansion velocity, depending of dispersal mortality (case where $r_0 = \log(1.5)$, see **main text Fig. 3** for the case where $r_0 = \log(5)$). Predictions are based on the best model **Fig. S5.1**; predictions for the effect of $d_0$ are made assuming no density-dependence ($\Delta_{avg-0} = 0$), and predictions for the effect of $\Delta_{avg-0}$ are made setting $d_0$ to its average value in the simulated dataset.

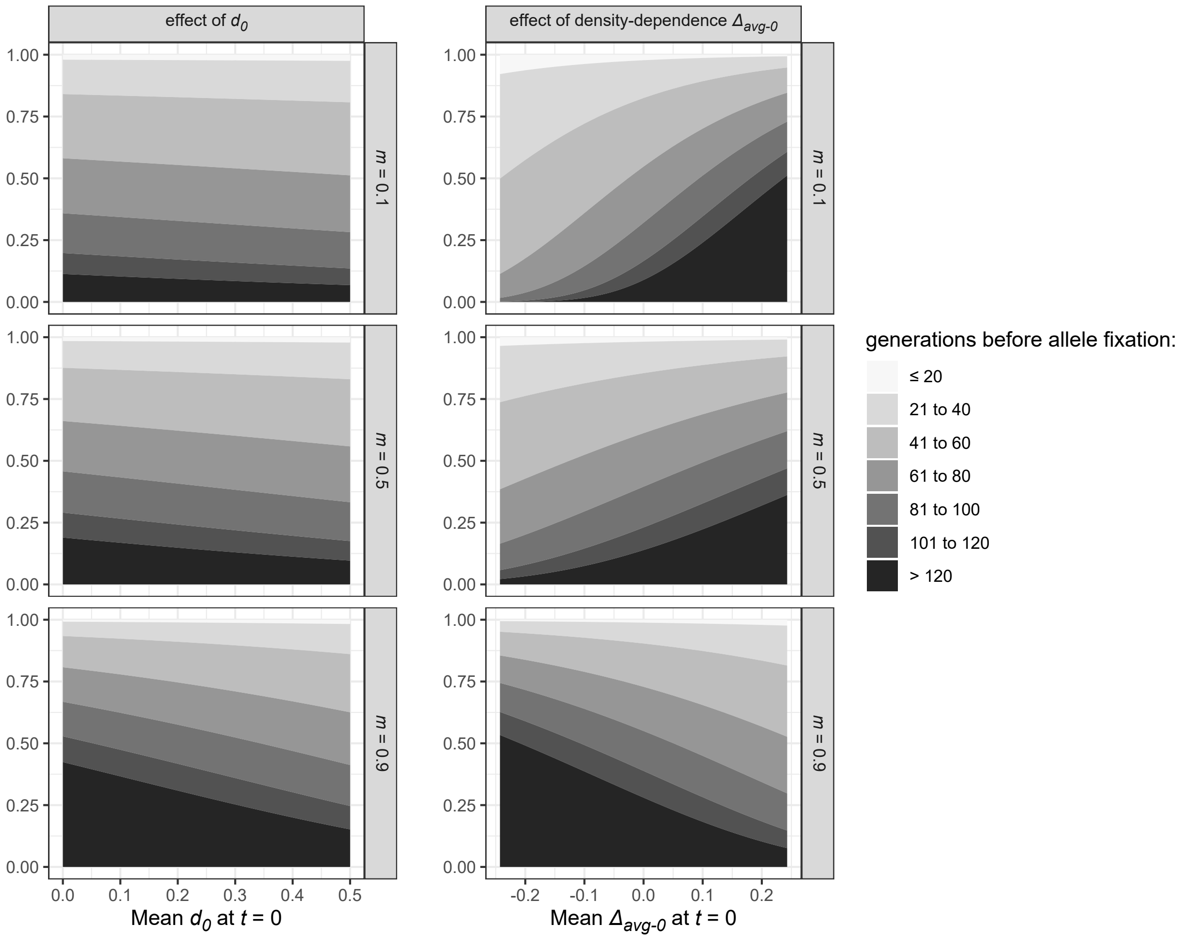


**Figure S5.4**. Predicted effect of $d_0$ (left) and density-dependence $\Delta_{avg-0}$ (right) on the loss of neutral diversity at the range front, depending of dispersal mortality (case where $r_0 = \log(1.5)$, see **main text Fig. 4** for the case where $r_0 = \log(5)$). Darker shades of grey indicate later fixation times. Predictions are based on the best model **Fig. S5.1**; predictions for the effect of $d_0$ are made assuming no density-dependence ($\Delta_{avg-0} = 0$), and predictions for the effect of $\Delta_{avg-0}$ are made setting $d_0$ to its average value in the simulated dataset.

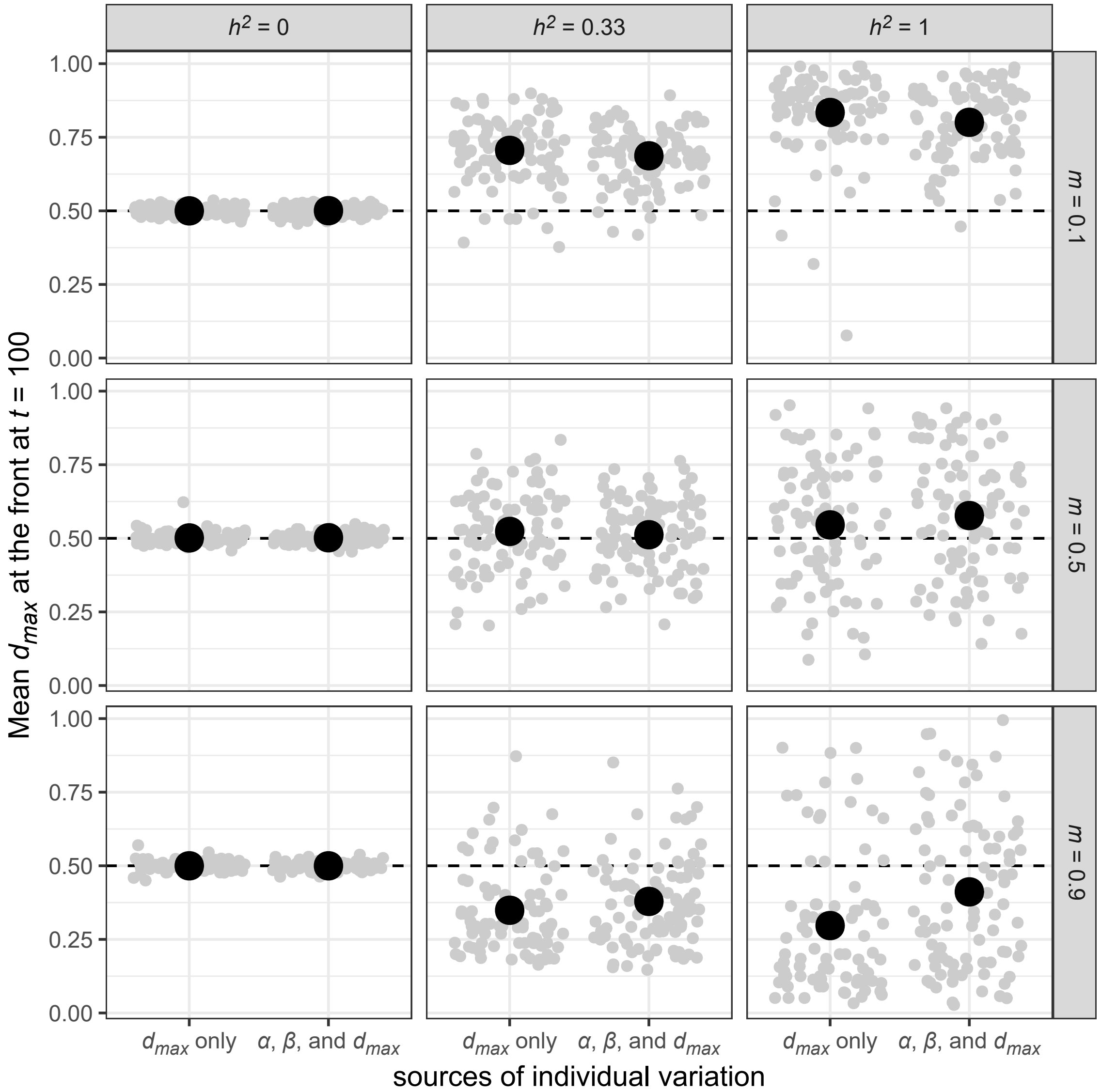


**Figure S5.5**. Evolutionary changes in mean dispersal capacity $d_{max}$ at the range front as a function of sources of phenotypic variation and dispersal costs (case where $r_0 = \log(1.5)$, see **main text Fig. 5** for the case where $r_0 = \log(5)$). Grey dots are individual replicates, with the average values overlaid in black. Replicates with no individual variation in $d_{max}$ at $t = 0$ are not displayed.

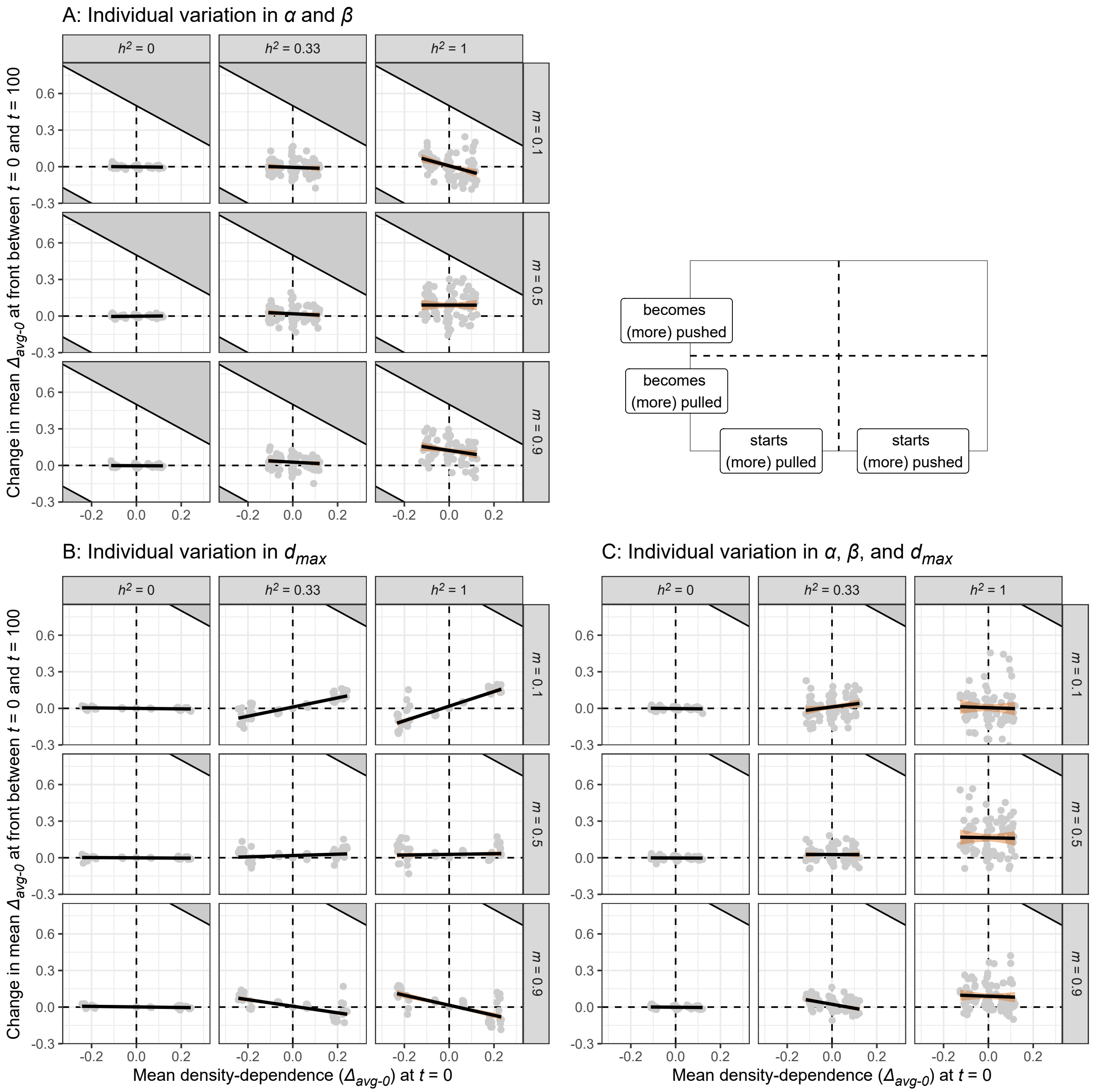


**Figure S5.6**. Evolutionary changes in mean density-dependence $\Delta_{avg-0}$ at the range front as a function of its initial value, sources of phenotypic variation and dispersal costs (case where $r_0 = \log(1.5)$, see **main text Fig. 6** for the case where $r_0 = \log(5)$). Grey dots are individual replicates, with linear regression lines overlaid. Replicates with no individual variation in traits are not displayed. Grey-filled areas indicate impossible phenotypes, more specifically evolved values of $\Delta_{avg-0}$ that would imply dispersal rates higher than the maximal possible $d_{max}$ (1 if $d_{max}$ is variable among individuals, 0.5 if it is not).

### S6 - An unconditionality metric

An interesting result from Travis et al. (2009) is that dispersal seems to become more density-independent at range edges. That is, the dispersal-density function becomes flat over a wider and wider range of densities.

However, we find (see main text) that density-dependence relevant to pushed vs pulled expansions still exists after evolution at the range edge. How to reconcile this?

It is relatively easy actually: the first step is to see that "flat over a larger and larger range of densities" does not mean "uniformly flat." As long as dispersal is slightly lower at specifically low densities, pushed dynamics can still happen, even if dispersal is fully independent of density over the rest of possible densities. The second step is to define an actual unconditionality index, and to test the prediction that, despite our results, our expansions still tend to become unconditional, on average.

For a given emigration-density function $f(n)$ and a relevant range of densities $[0 - K]$, we can define an unconditionality index $u$ as follows:

$$u = \frac{\int_0^K f(n)\mathrm{d}n}{\max\limits_{0 \leq n \leq K} f(n)}.$$

This index can take values between 0 and 1. The closer this index is to 1, the closer the average dispersal rate over the range 0-$K$ is equal to the maximal dispersal rate over the same range, i.e. the closer the dispersal function is to an horizontal line. For simplicity, as numerical integration from within base NetLogo is not possible, we approximated the numerator of $u$ by instead sampling $l$ density values at regular intervals over the range of densities and averaging the resulting dispersal rates:

$$N = \{0, 0.1K, 0.2K, ..., 0.9K, K\},$$

$$u_{[approx]} = \frac{(\sum_{n \in N} f(n))/l}{\max\limits_{n \in N} f(n)}.$$

We can then easily see that evolution of dispersal matching traits ($\alpha$ and $\beta$) indeed leads dispersal to become more unconditional when fecundity is high (**Fig. S6.1**).

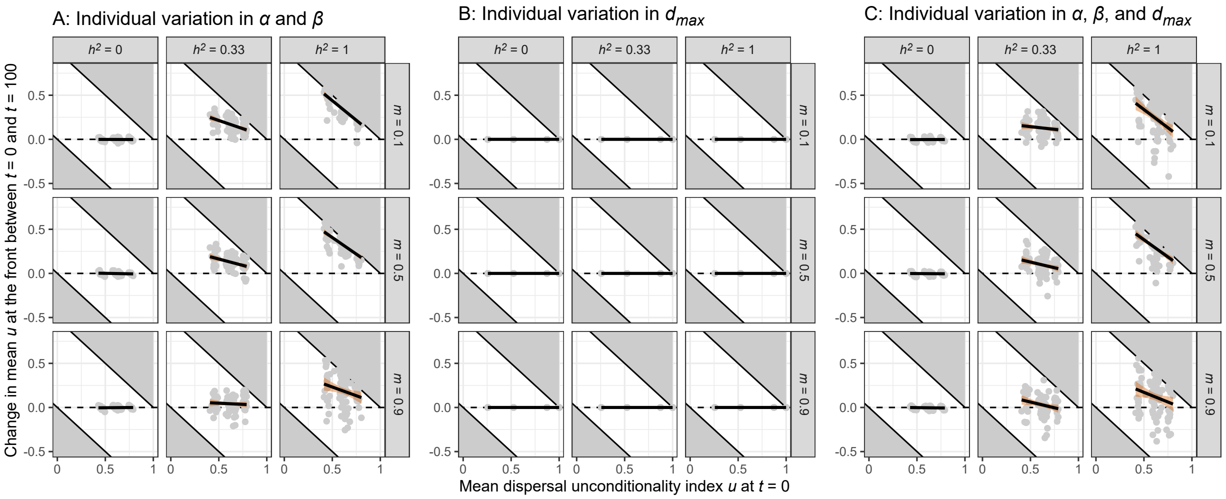


**Figure S6.1**. Evolutionary changes in mean dispersal unconditionality $u$ at the range front as a function of its initial value, sources of phenotypic variation and dispersal costs (case where $r_0 = \log(5)$, see **Supplementary Figure S6.2** for the case where $r_0 = \log(1.5)$). Grey dots are individual replicates, with linear regression lines overlaid. Replicates with no individual variation in traits are not displayed. Grey-filled areas indicate impossible states, i.e. changes in $u$ that would lead to $u > 1$ or $u < 0$.

When fecundity is reduced, dispersal still becomes more unconditional with expansion, unless mortality costs are high (**Fig. S6.2**, below). Jointly with our main text result, this confirms that metrics specifically targeting differences to low-density behaviour are needed to understand pushed expansions.

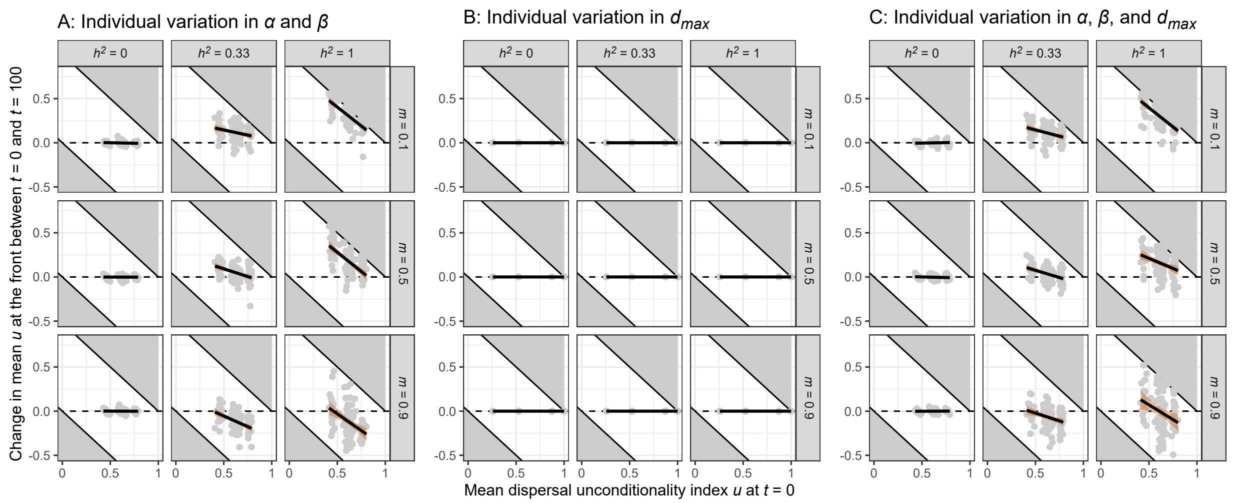


**Figure S6.2**. Evolutionary changes in mean dispersal unconditionality $u$ at the range front as a function of its initial value, sources of phenotypic variation and dispersal costs (case where $r_0 = \log(1.5)$, see **Supplementary Figure S6.1** for the case where $r_0 = \log(5)$). Grey dots are individual replicates, with linear regression lines overlaid. Replicates with no individual variation in traits are not displayed. Grey-filled areas indicate impossible states, i.e. changes in $u$ that would lead to $u > 1$ or $u < 0$.
